# Satellite glial contact enhances differentiation and maturation of human iPSC-derived sensory neurons

**DOI:** 10.1101/2024.07.24.604966

**Authors:** Chelsey J. LeBlang, Maria F. Pazyra-Murphy, Elizabeth S. Silagi, Srestha Dasgupta, Marianna Tsolias, Toussaint Miller, Veselina Petrova, Shannon Zhen, Vukasin Jovanovic, David Castellano, Kathryn Gerrish, Pinar Ormanoglu, Carlos Tristan, Ilyas Singeç, Clifford J. Woolf, Ozge Tasdemir-Yilmaz, Rosalind A. Segal

## Abstract

Sensory neurons generated from induced pluripotent stem cells (iSNs) are used to model human peripheral neuropathies, however current differentiation protocols produce sensory neurons with an embryonic phenotype. Peripheral glial cells contact sensory neurons early in development and contribute to formation of the canonical pseudounipolar morphology, but these signals are not encompassed in current iSN differentiation protocols. Here, we show that terminal differentiation of iSNs in co-culture with rodent Dorsal Root Ganglion satellite glia (rSG) advances their differentiation and maturation. Co-cultured iSNs develop a pseudounipolar morphology through contact with rSGs. This transition depends on semaphorin-plexin guidance cues and on glial gap junction signaling. In addition to morphological changes, iSNs terminally differentiated in co-culture exhibit enhanced spontaneous action potential firing, more mature gene expression, and increased susceptibility to paclitaxel induced axonal degeneration. Thus, iSNs differentiated in coculture with rSGs provide a better model for investigating human peripheral neuropathies.

## Introduction

Dorsal Root Ganglion (DRG) neurons are responsible for communicating somatosensory input from the outside world to the central nervous system^1^. They relay these signals through a single specialized T-shaped axon, which projects to both the periphery and the spinal cord^1–3^. DRG neurons include multiple phenotypic subtypes, which respond to specific stimuli and exhibit distinct patterns of gene expression^1,4,5^. DRG neurons are subject to a large number of disorders including hereditary neuropathies, diabetic neuropathy, and chemotherapy induced peripheral neuropathy^6^, therefore developing appropriate human model systems is a major goal for researchers.

The use of differentiated human induced pluripotent stem cells (hiPSCs) has become a gold standard for studying disease pathology in relevant cell types and is increasingly accessible^7,8^. Over the past decade, several protocols have been generated to differentiate standard and patient derived hiPSCs into somatosensory neurons (iSNs, hiPSC-derived sensory neurons)^9–15^. Researchers have developed these protocols and utilized iSNs to study neuropathic pain signaling^10,16–18^, mechanisms underlying debilitating and painful neuropathies^19–24^, and development of treatments for these disorders^17,23,25,26^.

During sensory neuron differentiation, iPSCs undergo changes mimicking embryonic development, altering their transcriptional program to progressively resemble neuroectoderm, neural crest progenitor cells, and ultimately sensory neurons, through treatment with combinations of small molecules over time and/or genetic manipulation^9–15^. Cells produced using these protocols express many hallmark proteins associated with sensory neuron identity, however iSNs exhibit an embryonic phenotype, different from that of postnatal neurons^4^. In the developing mammalian embryo, DRG neurons initially are bipolar in morphology and undergo a series of morphological changes leading to a pseudounipolar morphology before birth^27–29^. Current differentiation protocols generate iSNs that are predominantly bipolar in morphology, reflecting the early embryonic state, or a multipolar morphology, similar to that observed following an acute injury *in vivo*^30–32^. While developing sensory neurons alter their transcriptome over the course of prenatal development and express transcription factors that contribute to neuronal subtyping^1,33^, iSNs remain functionally unspecialized. Therefore, modifications of sensory neuron differentiation protocols to achieve a further differentiated phenotype would greatly advance efforts to study postnatal homeostasis and disease at a molecular level, in a way that is more translatable to human health or disease conditions.

We have sought to leverage developmental cues to advance the maturation of differentiated sensory neurons. While specific mechanisms underlying pseudounipolarization and subtyping are unknown, *in vitro* studies show that primary sensory neurons co-cultured with mixed DRG glia, including Schwann cells (SCs) and satellite glia (SGs), become pseudounipolar^34–36^. Yet, iSNs differentiated in a co-culture with induced or primary SCs, undergo no morphological changes, aside from myelination^16,37,38^. *In vivo,* intimate contact is initiated between sensory neurons and satellite glia during the early developmental period^39,40^, therefore we hypothesized that satellite glia contribute to DRG sensory neuron maturation, and that terminal differentiation of iSNs in coculture with satellite glial cells would advance this. Here, we show that physical contact between satellite glia and iSNs during differentiation significantly advances morphological and physiological development compared to those differentiated alone, and both semaphorin-plexin signaling and gap junction signaling contribute to this maturation process. We also tested the utility of this model and demonstrate that cocultured iSNs are more susceptible to toxic neuropathy compared to iSNs alone. This work not only provides an advanced iSN model for use by researchers in future studies, but also provides insight into mechanisms underlying sensory neuron development.

## Results

### Human iSNs differentiated in coculture with embryonic rodent DRG glia exhibit advanced morphology

Currently, differentiation protocols used to generate iSNs from iPSCs produce sensory neurons with predominantly immature morphologies^9–15^. While the molecular mechanisms underlying embryonic transition to pseudounipolar morphology are unknown, our data (Fig. S1A-B) and others^36,40^ suggest that satellite glia may play a pivotal role in morphological maturation, through contact with sensory neurons during this developmental period. To test this, we used our previously published protocol^15^ to terminally differentiate human iPSC-derived neural crest precursors (iNCPs) alone, or together with peripheral glia prepared from rat E14.5 DRGs (Fig. 1A). After one week, cells were fixed and stained to visualize neuronal morphology (TUJ1, cytoskeletal marker, green) and contact with rDRG satellite glia (p75, NGF receptor, magenta) (Fig 1B).

**Figure 1:**
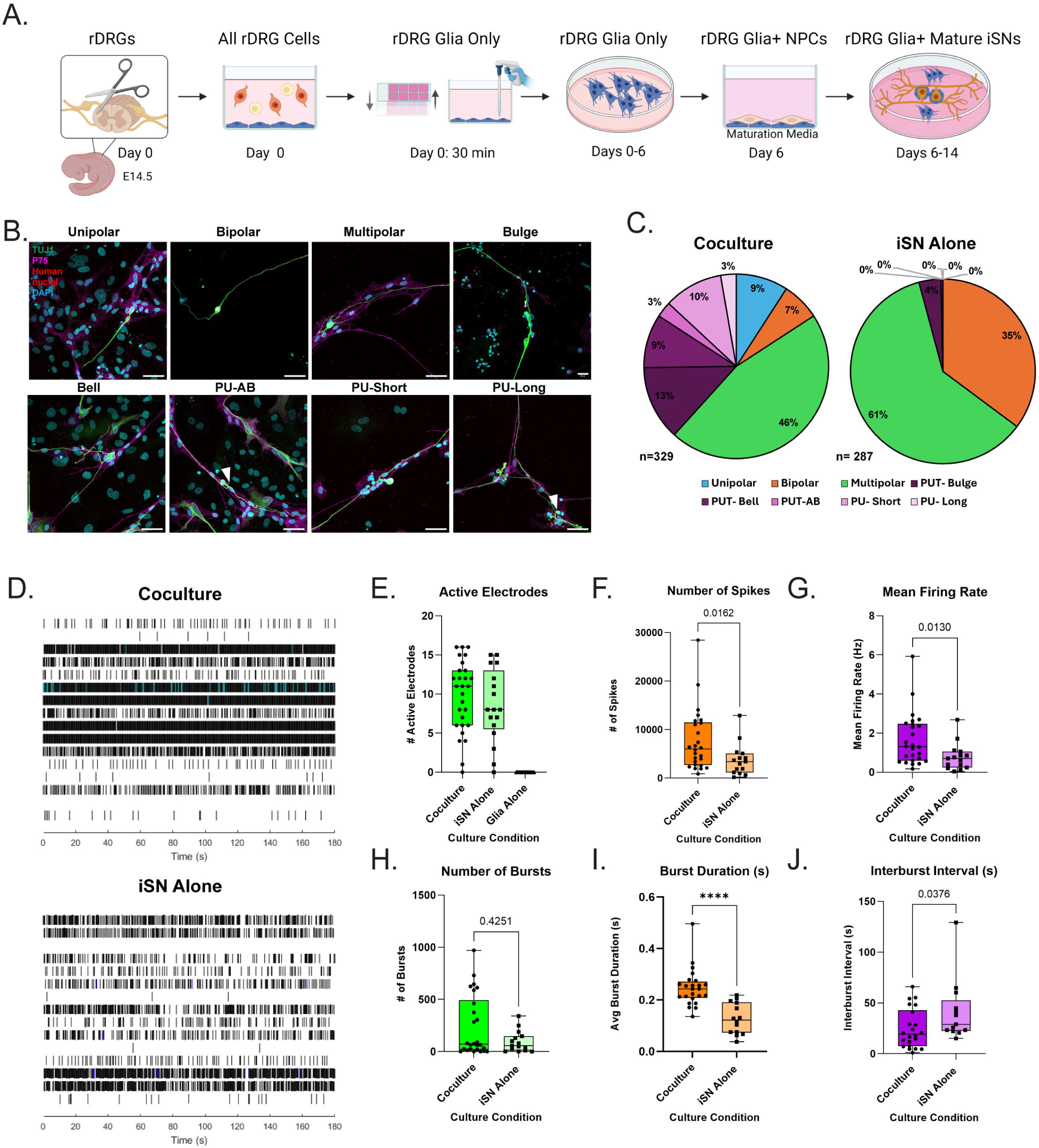
Human iSNs differentiated in coculture with embryonic rodent DRG glia exhibit more mature morphology and physiology. A) Schematic illustrating protocol to terminally differentiate iSNs in coculture with embryonic rat DRG glia (rDRG). Briefly, DRGs are dissected from E14.5 rat embryos. After 30 minutes, cultureware is shaken and cells in suspension are removed. Glia are cultured for 1 week, after which iNCPs are thawed and plated in coculture or alone for 1 week of maturation. B) Photomicrographs representing distinct iSN morphologies observed. TUJ1= Green, p75= Magenta, Human Nuclei= Red, DAPI= Cyan. Scale=50um. PU= pseudounipolar. PU-AB: white arrowhead indicates accessory branch. PU-Long: white arrowhead indicates the PU-Long neuron in this field. C) Frequency of each morphology observed in coculture and alone, displayed as percentages. 4 biological replicates, 2 iPSC cell lines. D. Representative raster plots over 2.5 minutes of MEA recording. Black lines indicate individual spikes and blue lines indicate bursts. E) Average number of active electrodes when average was > 7 (5-7 weeks). Coculture n= 28 wells, iSN alone n=17 wells, Glia alone n=16 wells from 4 biological replicates. F-H) Number of spikes (F), mean firing rate (G), and number of bursts (H) when average # active electrodes were >7. Wells with <2 active electrodes were excluded. Coculture n= 26 wells, iSN alone n=15 wells from 4 biological replicates. I) Average burst duration in seconds when average # active electrodes were >7. Wells with <2 active electrodes and 0 bursts were excluded. Coculture n=24 wells, iSN alone n=14 wells from 4 biological replicates. J) Interburst Interval in seconds when average # active electrodes were >7. Wells with <2 active electrodes and <2 bursts were excluded. Coculture n= 22 wells, iSN alone n=13 wells from 4 biological replicates. F-J) Mann-Whitney U tests performed in Graphpad Prism 10, p<0.05=significant, p<0.0001=****.

Morphologies of human neurons were identified and quantified using the rules specified in the methods section. Morphology categories compared consisted of “immature” (including unipolar and bipolar cells) “multipolar”, “Pseudounipolar Transition (PUT)” (including PUT-Bulge, PUT-Bell, and PU-Associated Branch), and “pseudounipolar (PU)” (including PU-Short and PU-Long) (Fig. 1B). Frequencies observed for each morphology category (Table 1) indicate that iSNs differentiated in coculture include significantly more transitioning and pseudounipolar cells (PUT *adj. residual* = 7.215, PU *adj. residual* = 6.350) and fewer cells with an immature morphology (immature *adj. residual* = -5.555, multipolar *adj. residual* = -3.653) compared to iSNs differentiated alone (PUT *adj. residual* = -7.215, PU *adj. residual* = -6.350, immature *adj. residual* = 5.555, multipolar *adj. residual* = 3.653) (Fig. 1C). No differences were observed between iSNs derived from different iPSC cell lines. These results suggest that terminal differentiation in co-culture with DRG glia enhances the maturation of sensory neuron morphology.

**Table 1.** Raw frequency of morphology subtypes observed in co-cultured iSNs vs. iSNs terminally differentiated alone. X^2^ frequency statistic calculated for major morphology groups: Immature, Multipolar PUT, and PU. (X^2^= 111, df=3, p<0.001).

|  | <u>Morphology Frequencies</u> |  |  |  |  |  |  |  |  |
| --- | --- | --- | --- | --- | --- | --- | --- | --- | --- |
|  | Immature |  | Multipolar | Pseudounipolar Transition (PUT) |  |  | Pseudounipolar (PU) |  |  |
| Condition | Unipolar | Bipolar | Multipolar | PUT-Bulge | PUT-Bell | PUT-AB | PU-Short | PU-Long | Totals |
| Coculture | 30 | 22 | 151 | 43 | 31 | 9 | 34 | 9 | 329 |
| iSN Alone | 0 | 101 | 174 | 11 | 1 | 0 | 0 | 0 | 287 |
| Totals | 153 |  | 325 | 95 |  |  | 43 |  | 616 |

To determine whether the ability of peripheral glia to promote a mature, pseudounipolar morphology is restricted to only one of the protocols used to generate iSNs, we also tested our co-culture protocol using RealDRG^TM^ neurons from Anatomic. RealDRGs were terminally differentiated in coculture or alone, using Senso-MM maturation media by Anatomic as a control. RealDRGs cocultured with rDRG Glia in NCATS Maturation Media (NMM)^15^ undergo extensive morphologic changes, although the frequency of transition to pseudounipolar morphology was reduced compared to Deng et al. 2023^15^ generated iSNs (Fig S1C-D). Treatment of RealDRGs cultures alone with NMM did not impact observed cell health or morphology but did result in the presence of neural precursor cells, which are not typically observed in RealDRG cultures maintained in Anatomic Senso-MM Media (Fig S1C-D).

### Human iSNs differentiated in coculture with embryonic rodent DRG glia exhibit advanced physiology

We next assessed the impact of the coculture strategy on development of spontaneous electrophysiologic signaling in iSNs. DRG sensory neurons can transmit sensory input to the CNS via single action potential spikes, as well as synchronous action potential bursts^41,42^. While spontaneous activity is relatively low^41–45^ increased spiking and bursting is observed in models of neuropathic pain both *in vivo*^41,42,45–48^ and *in vitro*^42,43,45,49^. Therefore, it is crucial that iSNs develop physiologic signaling capability to study the pathogenesis of neuropathies and develop potential treatments.

Characterization of action potential firing using Multi-Electrode Array (MEA) recordings shows that iSNs generated with the Deng et al. 2023^15^ protocol exhibit nearly undetectable baseline spontaneous activity after the typical two-week maturation period. Since peripheral satellite glial cells play an important role in the regulation of somatosensory neuron firing^50–53^, we tested the hypothesis that iSNs differentiated in coculture would exhibit more mature electrophysiologic properties by measuring spontaneous neural activity of iSNs differentiated in coculture, or alone, using Axion MEA plates containing 16 electrodes/well. Activity was measured on the Maestro Pro recording system for 5 minutes weekly over the span of 5-7 weeks, and neuronal spiking and bursting was analyzed using AxIS software. Overall, cocultured iSNs became robustly active earlier than iSNs differentiated alone, as indicated by a higher number of active electrodes beginning around week 3 of maturation (Fig. S1E-H). Data was analyzed when an average of 7 or more electrodes were active per well in both iSN culture conditions (Fig. 1E), this occurred between weeks 5-7 across 4 experimental replicates (Fig. S1E-G). Glia alone were inactive and do not contribute to signals recorded in the coculture (Fig. 1E, Fig. S1E-H).

Mature iSNs differentiated in coculture were more active than iSNs differentiated alone (Fig. 1E), with a significantly higher number of spikes elicited over the recording period (Mann-Whitney Test, p= 0.0162) (Fig. 1F), as well as a significantly higher Mean Firing Rate (MFR) (Mann-Whitney Test, p= 0.013) (Fig. 1G). The average MFR observed in cocultured iSNs, 1.657 Hz, is comparable to MFRs previously reported in primary DRG neurons^54,55^ and iSNs cultured long-term^56^. No significant difference was observed in the total number of bursts recorded, however the range was quite different, with a maximum burst number of 971 detected in coculture and only 341 in iSNs alone (Fig. 1H). Interestingly, cocultured iSNs elicited significantly longer burst trains (Mann-Whitney Test, p= <0.0001) (Fig. 1I), with a significantly shorter period between bursts (Mann-Whitney Test, p= 0.0376) (Fig. 1J), thus spending more time in a burst state. These results suggest that cocultured iSNs are more physiologically mature than iSNs cultured alone.

### iNCPs terminally differentiated in co-culture are advanced in neuronal differentiation

To identify the transcriptional programs that promote iSN maturation in coculture, we carried out RNA sequencing of iSNs terminally differentiated in coculture or alone. mRNA reads from cocultures were aligned to both the human and rat genome to distinguish mRNAs derived from hiSNs and rDRG glia respectively. PCA analysis confirmed that iSNs terminally differentiated in co-culture cluster separately from iSNs differentiated alone, illustrating transcriptional differences between these two groups (Fig. S2). Differential gene expression analysis yielded 476 significantly (p<0.05) upregulated genes in cocultured iSNs (Fig. 2A, Table S1). Genes implicated in posterior thoracic neural crest development^57,58^ (*HoxB9*, p adj.=6.922e^-15^) and proliferation^59^ (*ID3,* p adj.=1.306e^-09^) were among the top genes upregulated. Importantly, the most highly upregulated gene, *NFIX* (p adj.=1.025e^-16^), is associated with onset of neuronal differentiation and quiescence of neural stem cells ^60–62^ signaling a shift towards a more mature neuronal phenotype. Interestingly, *NFIX*, has also been shown to regulate neuroblast branching in adult neurogenesis^63^. Additional changes in expression of genes underlying neuronal morphogenesis include significant upregulation of *GPRIN3* (p adj.=6.437e^-12^) and *RAI14* (p adj.= 4.910e^-09^), proteins essential to axonal outgrowth^64,65^ and dendritic development^66,67^ respectively, through regulation of actin cytoskeletal structure. These changes illustrate potential transcriptional differences that underly the morphological maturation in cocultured iSNs.

**Figure 2.**
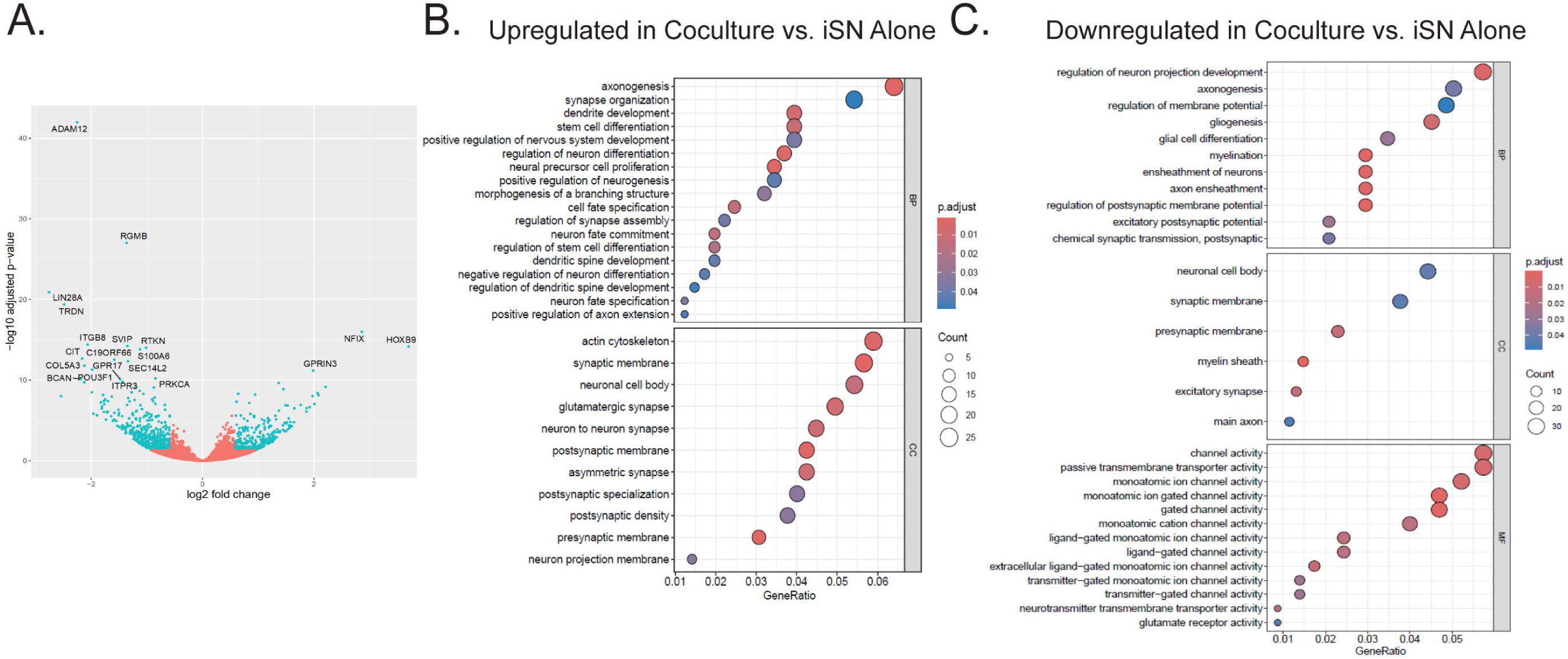
iNCPs terminally differentiated in co-culture more advanced in neuronal differentiation compared to iNCPs alone. A) Volcano plot illustrating differential gene expression in coculture vs. alone. B) GO analysis terms upregulated in coculture vs. iSNs alone, related to neuronal development. C) GO analysis terms downregulated in coculture vs. iSNs alone, related to neuronal development. Coculture n=3, iSN alone n=4. P adj.<0.05.

Gene Ontology (GO) analysis provided further information on the upregulated gene programs, with 185 significantly upregulated gene terms in cocultured iSNs compared to iSNs alone (Fig. 2B, Table S2). The most significantly upregulated biological processes include increased Bone Morphogenesis Pathway (BMP) signaling and anterior/posterior pattern specification, which confirm commitment to the neural crest cell lineage^33,68^ (Table S2). Several biological processes related to neuronal differentiation and morphogenesis were also significantly upregulated, including neural precursor proliferation, neuronal differentiation, and neuronal fate commitment, (Fig 2B). Interestingly, we also observed a significant upregulation of genes associated with axonogenesis (Fig 2B). These genes specifically influence axonal specification, outgrowth, guidance, branching, and synaptogenesis (Table 2, Table S2), and may potentially drive the pseudounipolarization process.

**Table 2.** Genes upregulated in cocultured iSNs that are implicated in aspects of axonogenesis.

| Role in Axonogenesis | Gene Names |
| --- | --- |
| Specification | *CDKL5*<sup>98</sup>/*PICALM*<sup>99</sup> |
| Outgrowth | *CDKL5*<sup>98,100</sup>/*AUTS2*<sup>101</sup> /*LAMA2*<sup>102</sup> /*PRICKLE1*<sup>103</sup>/ *EIF4G2*<sup>104</sup>/*TRPC5*<sup>105</sup> /*ADARB1*<sup>106</sup>/*ENAH*<sup>107</sup>/*RAB10*<sup>108</sup>, *USP9X*<sup>109</sup>/ *SOCS3*<sup>110</sup> |
| Branching | *NTN1*<sup>111</sup> / *BCL11A*<sup>112</sup> |
| Guidance | *EXT1*<sup>111</sup> /*PAK3*<sup>113</sup> /*EFNB2*<sup>114</sup>/ *NTN1*<sup>111,115</sup>/ *UNC5C*<sup>115</sup>, *GLI3*<sup>116</sup>, *FOXD1*<sup>117</sup> |
| Synaptogenesis | *NTN1*<sup>111</sup>/*SLITRK1/3*<sup>118</sup> /*ADARB1*<sup>119</sup> / *BCL11A*<sup>120</sup> / *DLG5*<sup>121</sup> |

Functional maturation was also evident with the upregulated expression of biological processes and cellular components related to synaptic structure and development, including synaptic membrane proteins, receptors, and ion channels localized to synapses in DRG neurons^4,69^ (Fig 2B). Specifically, we observed increased expression of genes encoding sodium ion channels (*SCN7a*/*SCN10a*) (Table S2)*. SCN10a* is highly expressed in DRG nociceptors, encoding the sodium channel Nav1.8^4,70^, critical for nociception^71–73^. We also observed upregulation of muscarinic receptors (*CHRM2/CHRM3*) essential to cholinergic signaling in DRG neurons^74–76^, and increased expression of potassium ion channels (*KCNA1/KCNQ5)* that limit action potential firing and maintain resting membrane potential in adult DRG sensory neurons^77–79^ (Table S2). Lastly, we observed significant upregulation in AMPAR gene *GRIA2* and GABA receptors (*GABRA1/GABRA5*) which are expressed in several adult DRG neuronal subtypes^4,69^ (Table S2). Overall, these transcriptional changes may underly the observed physiologic maturation of co-cultured iSN.

Conversely, 748 significantly (p<0.05) downregulated genes were identified in cocultured iSNs compared to iSNs alone (Fig. 2A, Table S1). There was decreased expression of genes implicated in early-stage neural crest development (*Lin28a*, p adj.=1.249^-21^; *RTKN,* p adj.=9.661^-15^)^80,81^ as well as a gene integral to differentiation of neural crest progenitors into mesenchymal lineage cells (*ADAM12*, p adj.=1.069^-42^)^82^ (Fig. 2A). We also observed a highly significant decrease in *RGMB* expression (p adj.=9.985^-28^), a BMP signaling partner which has been shown to promote ventral neuronal differentiation from neural crest cells^83^ (Fig. 2A). Additionally, we observed a significant decrease in expression of genes associated with peripheral gliogenesis, including myelinating Schwann cell markers (*Pou3F1*^84^, p adj.=4.872^-12^, *SVIP*^85^ p adj.=5.928^-15^, *GPR17*^86^ p adj.= 7.605^-1^*, MAL*^87^ p adj.=1.120^-8^*, PMP22*^88,89^ p adj.=1.325^-5^, satellite glial markers (*TRDN*^89^ p adj.=4.190^-20^, *BCAN*^88,89^ p adj.=7.605^-11^, *CHL1*^90^ p adj.=1.863^-7^), and glial proteins integral to axonal development (*S100A6*^91^ p adj.=1.481^-14^ *, SEMA3B*^92^ p adj.=8.410^-10^, *SEMA3G*^92^ p adj.=9.360^-8^) (Fig. 2A) suggesting enhanced commitment to a neuronal fate when iSNs are differentiated in co-culture.

Gene Ontology analysis yielded 134 significantly downregulated gene programs in iSNs differentiated in coculture compared to iSNs differentiated alone (Fig. 2C, Table S2). Gene programs associated with MHC I/II complex assembly and antigen presentation were among the most significantly downregulated biological processes in cocultured iSNs (Table S2), consistent with recent data that iNCPs express these proteins at low levels, if at all^93,94^. We also observed significant downregulation in gene programs associated with gliogenesis, glial differentiation, myelination, and axon ensheathment (Fig. 2C), consistent with results suggesting iSNs differentiated in coculture are committed to a neuronal, rather than a glial, fate. Interestingly, GO terms also included transcription factors implicated in the early specification of DRG neuronal subtypes including *NTRK3* and *SHOX2* (expressed in proprioceptors), *RET* (expressed in nonpeptidergic nociceptors and mechanoreceptors) *POU4f3* (expressed in a subpopulation of peptidergic nociceptors) and *GFRA3* (expressed in a subpopulation of peptidergic nociceptors)^1,4,95,96^ (Table S2). These results indicate that iSNs in co-culture are committed to a sensory neuron fate and restrict their transcription factor repertoire, which may advance development of functional DRG subtypes.

GO analysis also provides potential explanation for a change in neuronal physiology with decreased expression of genes involved in the development, composition, and function of synaptic membrane structures (Fig. 2C), including glutamate receptors, such as kainate receptors (*GRIK1/GRIK3*), AMPAR *GRIA4,* and Delta receptor *GRID2*^97^, as well as purinergic receptors (*P2RY1, P2RX6*) (Table S2). Interestingly, potassium ion channel expression (*KNCT2, KNCJ3, KCNJ12, KCNS1*) was also decreased, which could contribute to the increased excitability we observe in cocultured iSNs compared to iSNs differentiated alone^79^ (Table S2). Taken together transcriptomic analyses indicate that coculture of iNCPs with peripheral glia enhances differentiation into more mature sensory neurons.

### Rodent DRG satellite glia (rSGs) are responsible for advanced maturation of iSNs

Our studies demonstrate that iSNs differentiated in coculture with embryonic rat DRG cells mature at an expedited rate compared to iSNs differentiated alone, however due to the protocol used to prepare the rDRG cells, the glial subtype(s) driving this maturation was not initially clear. The rodent DRG cell population includes satellite glia (SGs), myelinating and non-myelinating Schwann cells (SCs), glial progenitor cells (GPCs), fibroblasts, endothelial cells, and immune cells. When initially plated (Fig. 1A, Day 0-7), rDRG cells display a multitude of morphologies, with small islands of semilunar cells and large flat cells, and this variety in cellular profiles is maintained as long as the cells remain in Glia Base Media (Fig. S3A-B). However, the morphology and cell number of rDRG glia changes upon addition of Deng et al. 2023^15^ NCATS Maturation Media (NMM), which is required for terminal differentiation of iSNs (Fig S3A-B). We used RNA sequencing to determine the impact of both NMM treatment and iSN contact on rodent glial identity and function (Fig. S3C), mRNA reads from co-cultured RNAs were aligned to both the human and rat genome to differentiate mRNAs derived from hiSNs and rDRG glia respectively.

PCA analysis and hierarchal clustering (Fig. S3D-E) showed that rDRG glia treated with NMM cluster together with cocultured rDRG glia treated with NMM. All glia treated with NMM cluster separately from rDRG Glia Alone maintained in base media, suggesting that media composition drives the transcriptional changes observed, while iSN contact leads to minimal changes in the glial transcriptome.

DGE analysis yielded 1160 significantly (p<0.05) upregulated genes in NMM treated rDRG glia compared to rDRG glia in Glia media (Fig 3A, Table S3). Overall, top upregulated genes in NMM treated glia are associated with mature satellite glia (SGs). The most upregulated genes, *DDIT4* (p adj.=1.196^-39^) and *INSIG1* (p adj.=1.196^-39^), have previously been identified as DRG SG specific markers in mouse sc-RNAseq studies^88,90^. *DDIT4* has been implicated in suppression of myelination^122^ and *INSIG1*^90^ plays a role in cholesterol biosynthesis. Lipid synthesis and metabolism are established functions of DRG SGs^123,124^ and all top upregulated genes with NMM treatment are involved in these processes including *PRR5*^125^ (p adj.=2.185^-23^), *PCSK9*^126^ (p adj.=2.067^-21^), *HMGCS1*^89^ (p adj.=5.928^-21^), *TMEM97*^127^ (p adj.=9.604^-19^). and *SCD*^89,90^ (p adj.=2.762^-18^). *HMGCS1* and *SCD*^89^, and their related upregulated genes, *HMGCR* (p adj.=1.537^-12^) and *SCD2*^90^ (p adj.=4.821^-15^), have also previously been identified as DRG SG gene makers in sc-RNA seq studies. GO analysis (Fig. 3B, Table S4) confirmed that lipid biosynthesis and metabolism pathways were significantly upregulated in NMM cultures.

**Figure 3.**
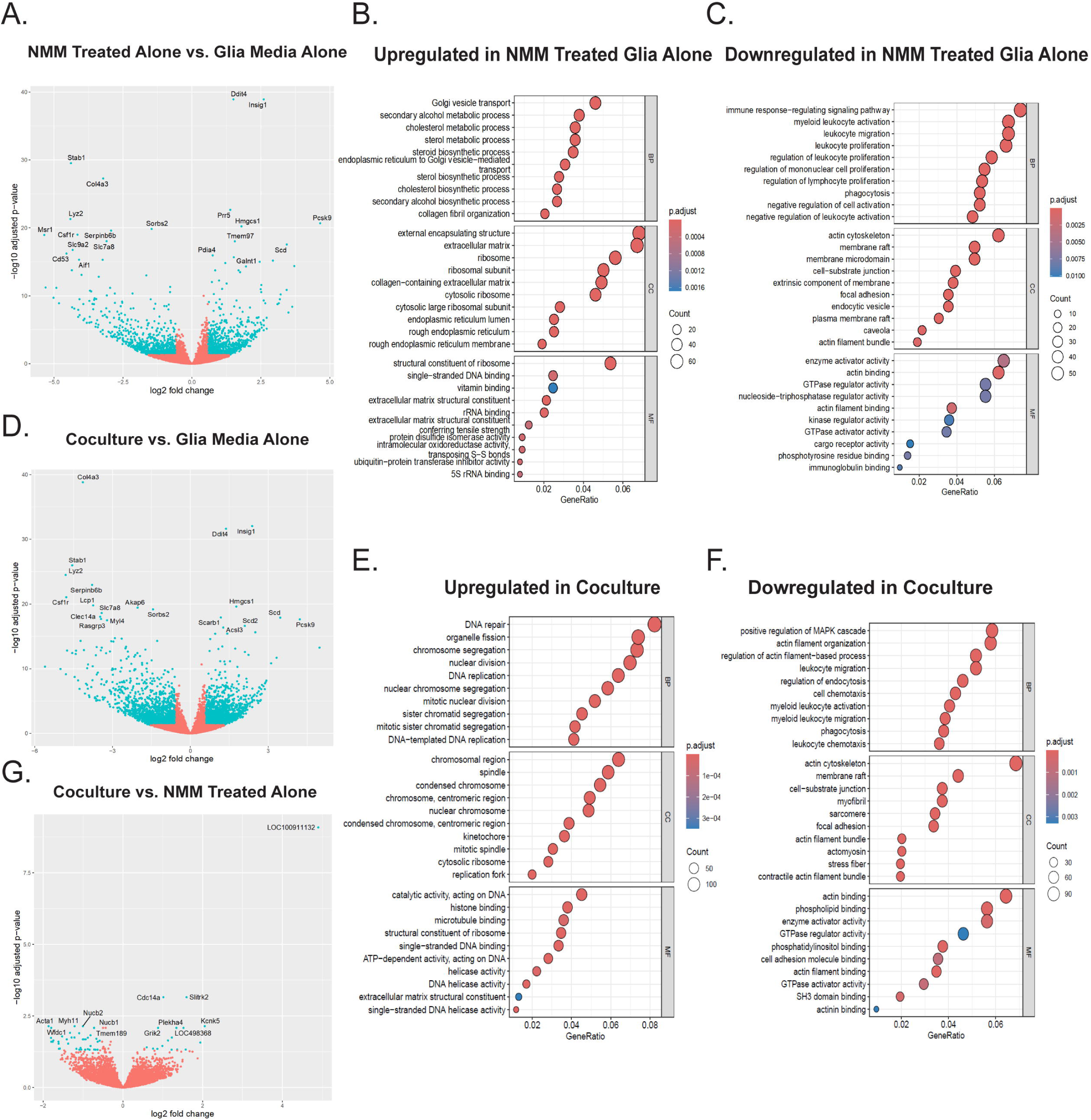
Rodent DRG satellite glia (rSGs) are responsible for advanced maturation of iSNs. A) Volcano plot illustrating differential gene expression in NCATS Maturation Media treated rDRG glia vs. rDRG glia maintained in base glia media B-C) GO analysis terms upregulated (B) and downregulated (C) in NCATS Maturation Media treated rDRG glia vs. rDRG glia maintained in base glia media. D) Volcano plot illustrating differential gene expression in cocultured rDRG glia vs. rDRG glia maintained in base glia media E-F) GO analysis terms upregulated (E) and downregulated (F) in cocultured rDRG glia vs. rDRG glia maintained in base glia media. G) Volcano plot illustrating differential gene expression in cocultured rDRG glia vs. NCATS Maturation Media treated rDRG glia. n=3-4/group. P adj.<0.05.

We also observed significant upregulation in biologic processes, cellular components, and molecular functions related to extracellular matrix (ECM) production. These include increased expression of SG genes related to collagen production, such as *COL16A1*, *COL1A1, COL3A1,* and *COL5A1*^90^, and other ECM related genes previously associated with SG identity, *FBLN5*^89,90^ and *ANXA5*^90^ (Table S4). Significant upregulation in biological processes related to DNA replication (Table S4) indicate that enrichment for SGs may be due to enhanced cellular proliferation. Together these results support that treatment with NMM results in an increased proportion of mature SGs in our cultures.

DGE analysis also yielded 933 significantly (p<0.05) downregulated genes in NMM treated rDRG glia compared cultures maintained in base Glia Media (Fig. 3A, Table S3). Overall, these results indicate a decrease in immune functions in NMM treated cultures. Top downregulated genes were key markers of DRG macrophages including *STAB1*^128,129^ (p adj.=2.984^-30^), *LYZ2*^89^ (p adj.=4.974^-22^), *CSF1R*^130^ (p adj.=9.916^-20^), *MSR1*^131^ (p adj.=1.090^-19^), and *AIF1*^89^ (p adj.=4.859^-16^). GO analysis (Fig. 3C, Table S4) confirmed that the most downregulated biological processes in NMM treated rDRG glia are related to various aspects of leukocyte function including leukocyte differentiation, proliferation, homeostasis, activation, migration, cytokine production, and chemotaxis. Significant decrease in expression of gene programs related to both mononuclear cells (phagocytosis, macrophage activation, macrophage differentiation) and lymphocytes (T-cell proliferation, T-cell activation, B-cell differentiation, B-cell activation, adaptive immune response) are observed (Table S4). GO analysis also reveals decreased expression of genes related to several aspects of actin cytoskeletal organization, which may contribute to morphologic changes observed with NMM treatment. Together these data indicate the composition of NMM results in an enrichment of SGs. When we compared transcriptomic differences between co-cultured rDRG glia treated with NMM and rDRG glia maintained in Glia Media, we found that differentially expressed genes were nearly identical to those identified in NMM treated cultures without iSN contact (Fig. 3D-F, Table S5-6), indicating that NMM treatment drives an enrichment of SGs in rDRG glial cultures.

To elucidate the impact of iSN contact on the transcriptome of SGs, we next compared gene expression between co-cultured NMM treated glia and NMM treated glia alone. DGE analysis yielded very few significant (p<0.05) differences, with only 20 significantly upregulated and 52 significantly downregulated genes in co-cultured NMM treated glia (Fig. 3G, Table S7). The results indicate that iSN contact impacts the SG cell-cycle and morphology with upregulation of genes related to DNA replication and repair (*CDC14A*^132^, p adj.=0.0007), transcription (*ZSCAN18,* p adj.= 0.0352; *TLE4* p adj.= 0.0393), translation (*QARS*^133^, p adj.=8.165^-10^), cytoskeletal organization (*DNMBP*^134^ p adj.= 0.047, *SRCIN1*^135^ p adj.= 0.047)*, SLITRK2*^136^ p adj.=0.0007) and ion channels (*TRPM3* p adj.= 0.047*, GRIK2* p adj.= 0.008*, KCNK5* p adj.= 0.007). Several upregulated genes were previously linked to specialized SGs (*SLITRK2*^86^*, TRPM3*^137^*, ITGB4*^90^*, GBP2*^90,137^), which may indicate changes in SG specific functional phenotype when in contact with iSNs. Conversely, downregulated genes (Fig. 3G, Fig. S3F, Table S7-8) were associated with collagen production (*P4HA3*^138^ p adj.= 0.013, *COL4A5*^139^ p adj.= 0.015) and cell-adhesion (*LTBP2*^140^ p adj.= 0.008), which may indicate an alteration in cell spreading and migration with neuronal contact.

Taken together, transcriptomic analyses illustrate that initial DRG cell preparations contain multiple cell types, however maturation media enriches for satellite glia. SGs in coculture and SGs cultured alone with NMM treatment are transcriptionally very similar, however minor changes in morphological and functional aspects of SGs do occur with neuronal contact.

### Rodent SG (rSG) secreted factors alone do not induce a pseudounipolar morphology in iSNs

The transcriptomic analysis confirmed that rodent satellite glia (rSGs) contribute to the maturation of iSNs, however the molecular mechanisms underlying this process are unknown. We first asked whether this process is mediated by glial secreted factors, using a transwell co-culture system (Fig. 4A). rSGs were cultured for one week on polycarbonate membranes inserted into a well plate, and iSNs were then terminally differentiated in the same wells beneath the membranes, sharing media but never physically touching rSGs, or in wells alone. iSNs were then fixed and stained to visualize neuronal morphology (TUJ1, cytoskeletal marker, green; BRN3a, iSN maturation marker, red) (Fig. 4B), while the rSGs on membranes were dyed and visualized live after two weeks of culture to confirm that a healthy population was present. iSNs differentiated in transwell coculture and alone both presented with predominantly multipolar or immature morphologies, with no significant transition to pseudounipolar morphology (Fig. 4C, Table 3).

**Figure 4.**
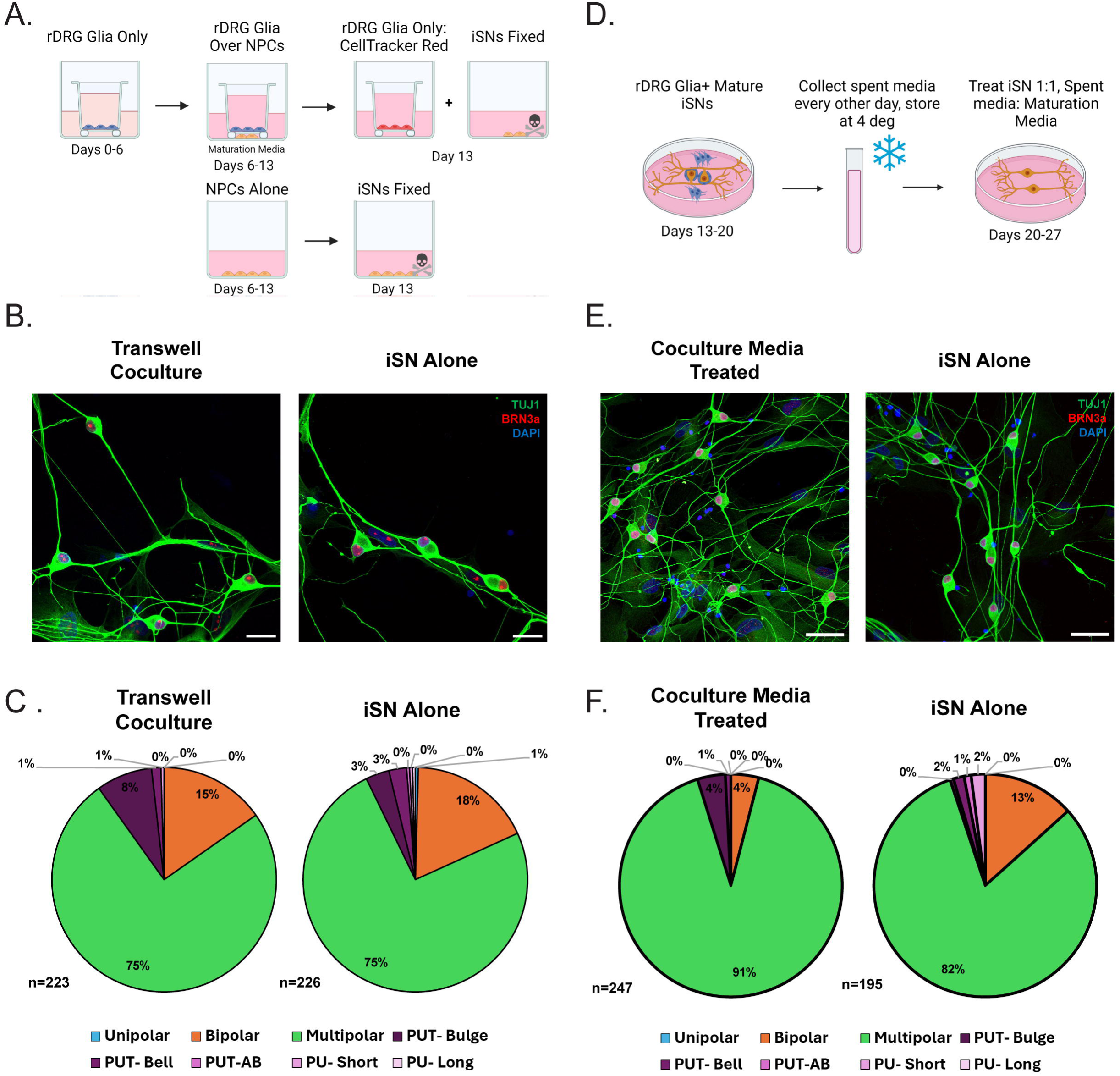
rSG secreted factors do not induce pseudounipolar morphology in iSNs. A) Schematic illustrating transwell coculture protocol. B) Representative photomicrographs of iSNs cultured in transwell coculture or alone. TUJ1= Green, BRN3a=Red, DAPI= Blue. Scale= 50um. C) Frequency of each morphology observed in coculture and alone, displayed as percentages .n= 3 biological replicates D) Schematic illustrating protocol for iSN treatment with coculture spent media. E) Representative photomicrographs of iSNs treated with coculture spent media or untreated. TUJ1= Green, BRN3a=Red, DAPI= Blue. Scale=50um. F) Frequency of each morphology observed in Coculture Media treated and untreated iSNs, displayed as percentages. 3 biological replicates.

**Table 3.**
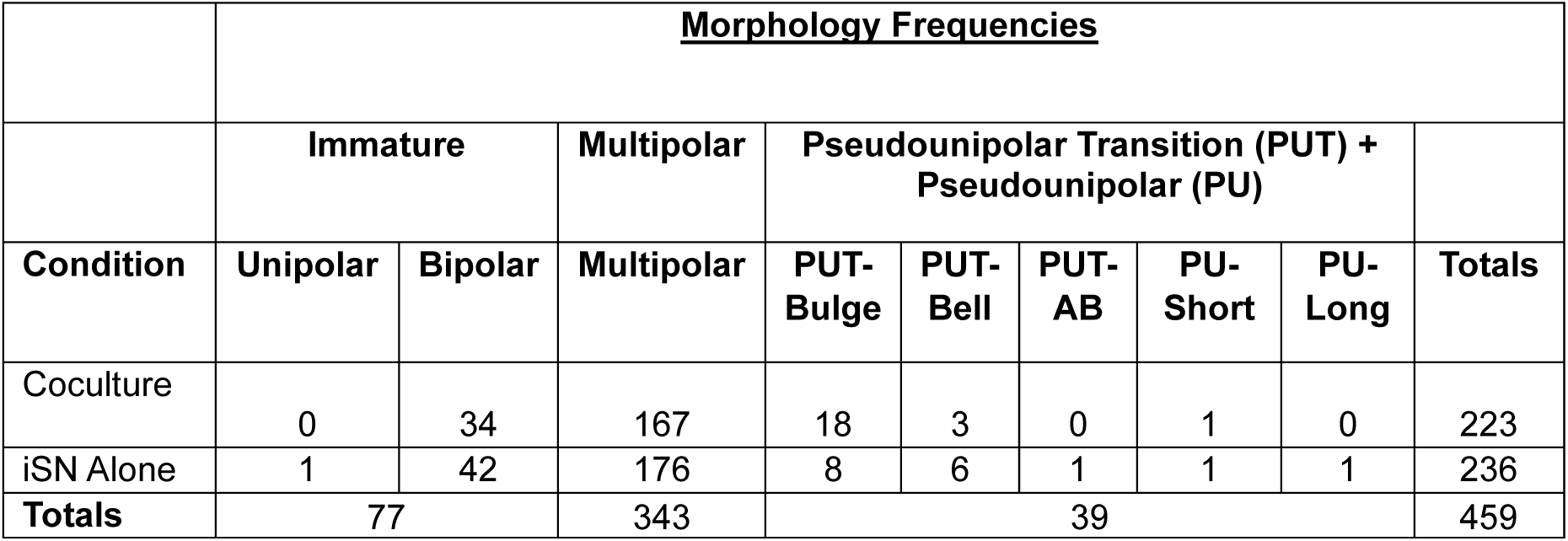
Raw frequency of morphology subtypes observed in transwell co-cultured iSNs vs. iSNs terminally differentiated alone. X^2^ frequency statistic calculated for major morphology groups: Immature, Multipolar, and PU(T). X^2^= 1.562, df=2, p= 0.4579.

|  | **<u>Morphology Frequencies</u>** |  |  |  |  |  |  |  |  |
| --- | --- | --- | --- | --- | --- | --- | --- | --- | --- |
|  | **Immature** |  | **Multipolar** | **Pseudounipolar Transition (PUT) + Pseudounipolar (PU)** |  |  |  |  |  |
| **Condition** | **Unipolar** | **Bipolar** | **Multipolar** | **PUT-Bulge** | **PUT-Bell** | **PUT-AB** | **PU-Short** | **PU-Long** | **Totals** |
| Coculture | 0 | 34 | 167 | 18 | 3 | 0 | 1 | 0 | 223 |
| iSN Alone | 1 | 42 | 176 | 8 | 6 | 1 | 1 | 1 | 236 |
| **Totals** | 77 |  | 343 | 39 |  |  |  |  | 459 |

It is possible that glia only secrete factors critical for the maturation process after they have made physical contact with iSNs. To test this possibility, iSNs plated alone were treated for six days with spent media collected from cocultures (Fig. 4D-F, Table 4). While there are no differences in pseudounipolar morphology, there is a higher frequency of immature neurons in untreated iSNs (immature *adj. residual* = 3.544) compared to spent media treated iSNs (immature *adj. residual* = -3.544), as well as an increased proportion of multipolar cells in spent media treated iSNs (multipolar *adj. residual* = 2.954) compared to untreated iSNs (multipolar *adj. residual* = - 2.954). These results are consistent with the idea that rSG-iSN contact, rather than secreted factors, are needed to induce pseudounipolar morphology.

**Table 4.**
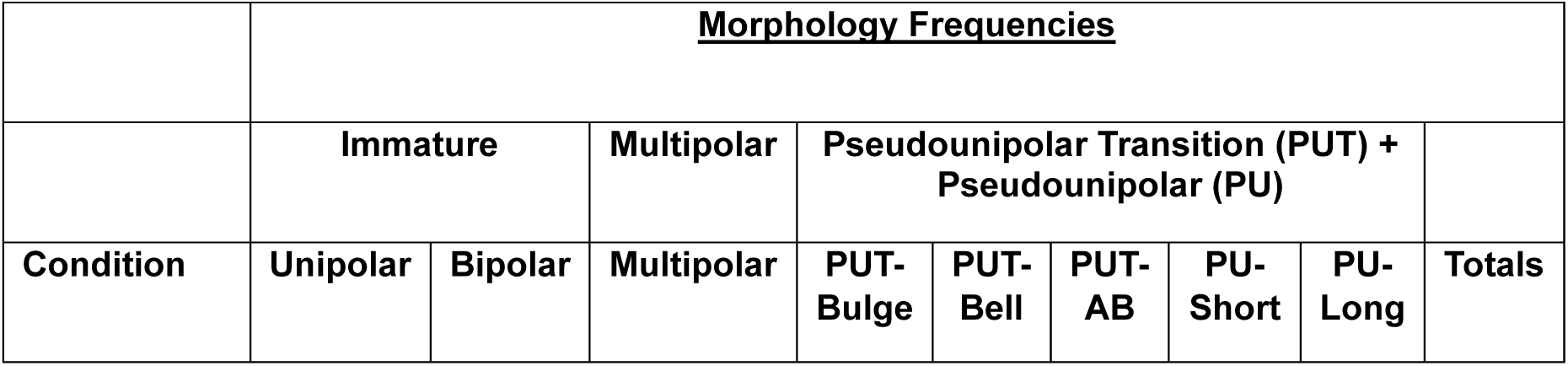
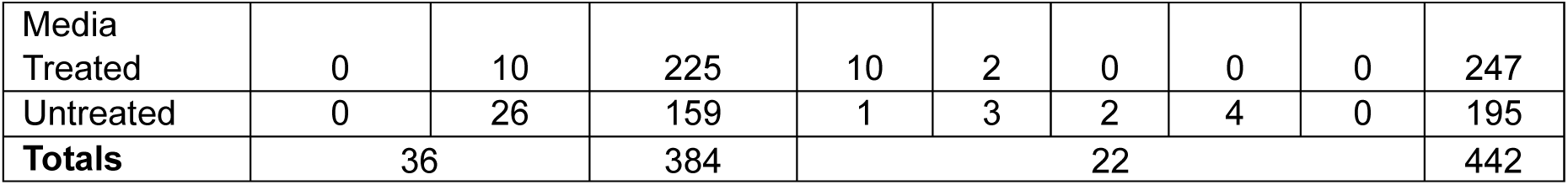
Raw frequency of morphology subtypes observed with and without coculture spent media treatment. X^2^ frequency statistic calculated for major morphology groups: Immature, Multipolar, and PU(T). X2= 12.69, df=2, p= 0.0018.

|  | **<u>Morphology Frequencies</u>** |  |  |  |  |  |  |  |  |
| --- | --- | --- | --- | --- | --- | --- | --- | --- | --- |
|  | **Immature** |  | **Multipolar** | **Pseudounipolar Transition (PUT) + Pseudounipolar (PU)** |  |  |  |  |  |
| **Condition** | **Unipolar** | **Bipolar** | **Multipolar** | **PUT-Bulge** | **PUT-Bell** | **PUT-AB** | **PU-Short** | **PU-Long** | **Totals** |
| Media Treated | 0 | 10 | 225 | 10 | 2 | 0 | 0 | 0 | 247 |
| Untreated | 0 | 26 | 159 | 1 | 3 | 2 | 4 | 0 | 195 |
| **Totals** | 36 |  | 384 | 22 |  |  |  |  | 442 |

### Physical contact between rSGs and hiSNs promotes pseudounipolarization

To determine whether physical contact with satellite glia promotes pseudounipolarization, we designed an assay in which rDRG cells were cultured for two weeks in NMM, then fixed with methanol prior to coculture, preserving surface protein configurations. iSNs were subsequently cultured with the methanol fixed glial cells, in a normal coculture, or alone for one week (Fig. 5A). Cultures were then fixed and stained to visualize neuronal morphology (TUJ1, cytoskeletal marker, green) and contact with rSGs (p75, NGF receptor, magenta) (Fig. 5B). iSNs differentiated in methanol fixed coculture exhibit a phenotype intermediate between iSNs differentiated in normal coculture and those differentiated alone (Fig. 5C, Table 5). The Chi-square frequency test identified significant differences between observed and expected morphology frequencies across all three groups (X^2^=131.3, df=6, p<0.001). Interestingly, these results are driven by significant differences between iSNs differentiated in normal coculture conditions (Immature *adj. residual* = -3.543, Multipolar *adj. residual*=-6.471, PUT *adj. residual*= 6.468, PU *adj. residual*=7.893) and iSNs differentiated alone (Immature *adj. residual* = 2.041, Multipolar *adj. residual* = 4.963, PUT *adj. residual*= -5.753, PU *adj. residual*= -3.893), while iSNs differentiated in a methanol fixed coculture exhibited “expected” morphology frequencies, with only significantly fewer pseudounipolar cells than expected (PU *adj. residual*= -4.007). Since PU(T) cells make-up 14.02% of iSNs differentiated in methanol fixed coculture, compared to the 2.26% of iSNs differentiated alone, these results suggest that physical contact between rSGs and iSNs contributes to the changes required for a transition to pseudounipolar morphology.

**Figure 5.**
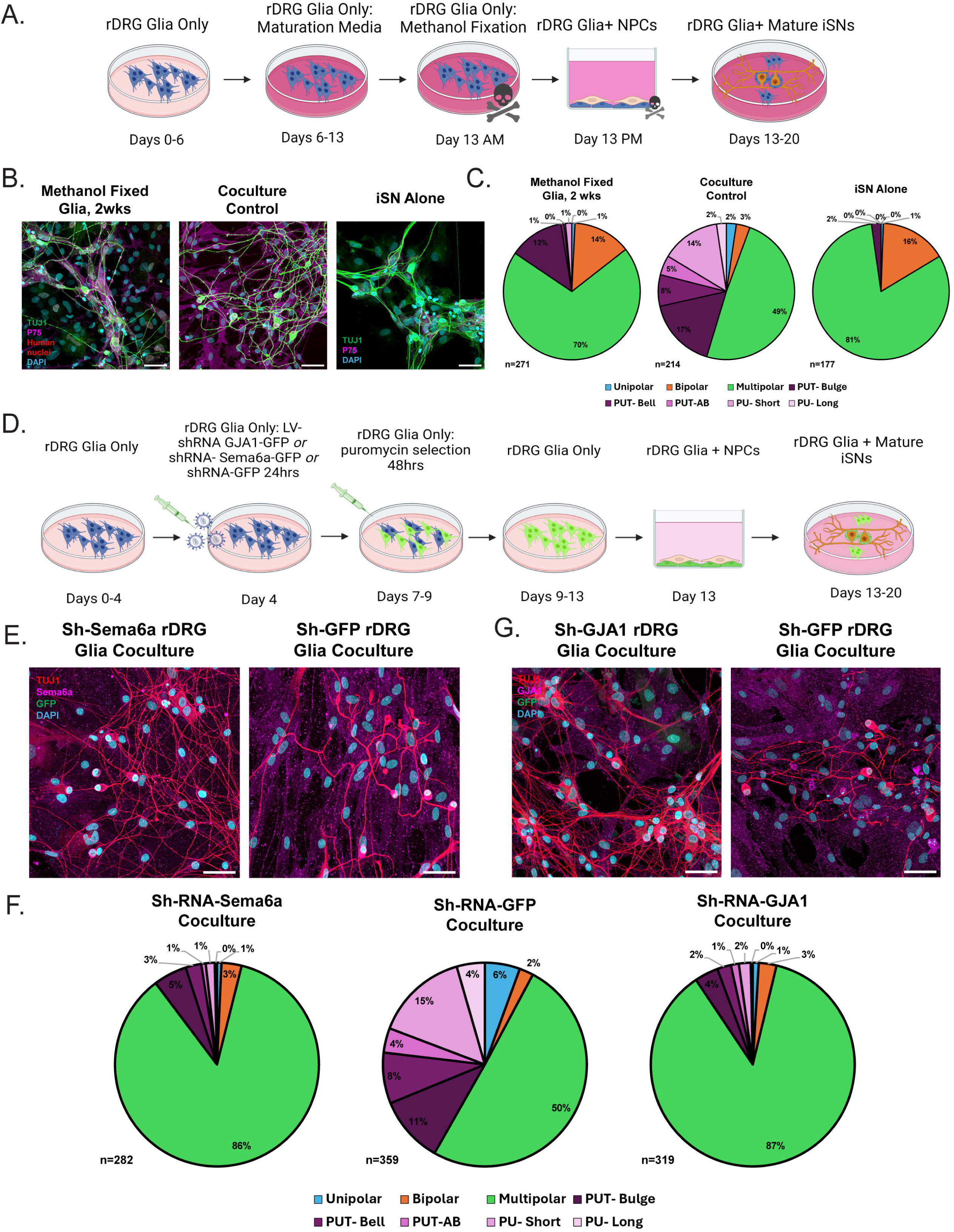
Physical contact between rSGs and iSNs is essential to pseudounipolarization. A) Schematic illustrating protocol for iSN coculture with methanol fixed glia. B) Representative photomicrographs of iSNs in coculture with methanol fixed glia, normal glia, or alone. TUJ1= Green, p75=Magenta, Human Nuclei= Red, DAPI= Cyan. iSNs alone were not stained for human nuclei. Scale= 50um. C) Frequency of each morphology observed in cocultured iSNs with methanol fixed glia and normal glia, displayed as percentages. n=4 biological replicates. D) Schematic illustrating protocol for iSN coculture with Sema6a, GJA1, and GFP knockdown in glia. All three conditions were run in parallel. E) Representative photomicrographs of iSNs in coculture with SEMA6a knockdown glia or GFP knockdown glia. Sema6a=Magenta, TUJ1= Red, GFP=Green, DAPI=Cyan. Scale=50um. F) Frequency of each morphology observed in iSNs cocultured with SEMA6a knockdown, GFP knockdown, and GJA1 Knockdown glia, displayed as percentages. n=4 biological replicates. G) Representative photomicrographs of iSNs in coculture with GJA1 knockdown glia or GFP knockdown glia. GJA1=Magenta, TUJ1= Red, GFP=Green, DAPI=Cyan. Scale=50um.

**Table 5:**
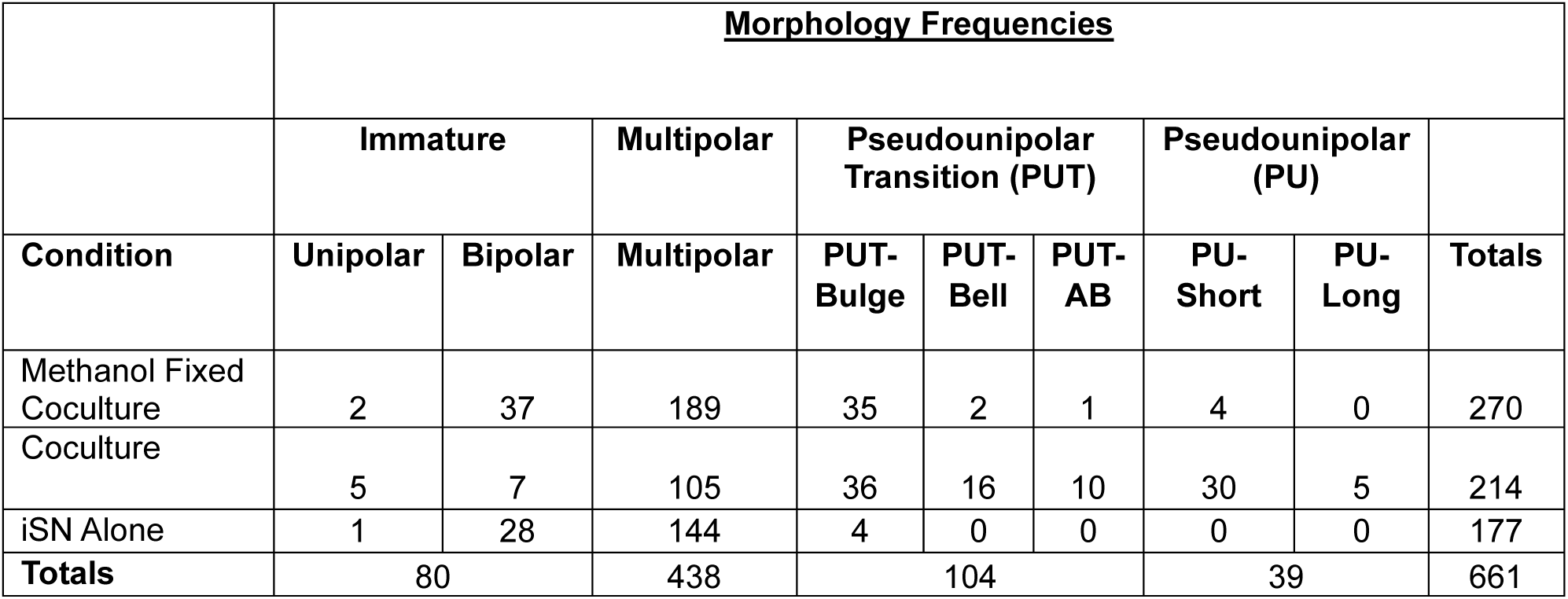
Raw frequencies of morphologies and morphology subtypes observed in methanol fixed coculture, normal coculture, and alone. X^2^ frequency statistic calculated for major morphology groups: Immature, Multipolar, PUT and PU. X2=131.3, df=6, p<0.001.

|  | <u>Morphology Frequencies</u> |  |  |  |  |  |  |  |  |
| --- | --- | --- | --- | --- | --- | --- | --- | --- | --- |
|  | Immature |  | Multipolar | Pseudounipolar Transition (PUT) |  |  | Pseudounipolar (PU) |  |  |
| Condition | Unipolar | Bipolar | Multipolar | PUT-Bulge | PUT-Bell | PUT-AB | PU-Short | PU-Long | Totals |
| Methanol Fixed Coculture | 2 | 37 | 189 | 35 | 2 | 1 | 4 | 0 | 270 |
| Coculture | 5 | 7 | 105 | 36 | 16 | 10 | 30 | 5 | 214 |
| iSN Alone | 1 | 28 | 144 | 4 | 0 | 0 | 0 | 0 | 177 |
| Totals | 80 |  | 438 | 104 |  |  | 39 |  | 661 |

The degree to which such direct contact truly contributes to this process is unclear, as methanol fixation prevents movement of rSGs in culture, and therefore physical contacts between rSGs and iSNs may be reduced compared to normal cocultures. Overall, the average number of rSG contacts per iSN was significantly reduced in the methanol fixed condition compared to normal cocultures (*t*=23.04, df=470, p<0.0001), with an average of 2 contacts/iSN in methanol fixed coculture vs. 10 contacts/iSN in normal cocultures (Fig. S4A). Significantly more contacts were made to true pseudounipolar cells in methanol fixed coculture compared to less advanced morphologies (Bonferroni post-hoc: PU vs. Multipolar p=0.0038; Bipolar p=0.0053, PUT Bulge p=0.0097) (Fig. S4B). In fact, pseudounipolar iSNs in methanol fixed co-culture were contacted to a similar degree to those in normal coculture, with an average of 9 rSG contacts/ iSN in methanol fixed coculture and 10 rSG contacts/iSN in normal coculture. These results indicate that contact is essential to pseudounipolarization, but downstream signaling and movements of live rSGs may also contribute to this process.

### Semaphorin-plexin and gap junction signaling play a role in pseudounipolarization of iSNs

We identified two potential pathways whose constituents are appropriately expressed to enable contact mediated SG signaling to iSNs, namely, Sema6a-Plexin A4 and gap junctions. Sema6a, a transmembrane semaphorin protein that is expressed in satellite glia, repels or inhibits axonal growth in developing sensory neurons through engagement with the PlexinA4 receptor^141–144^, which is highly expressed in hiSNs (Fig. S4C). Additionally, our sc-RNA seq data set from embryonic DRG satellite glia at the time of pseudounipolarization reveals high expression of *GJA1*, encoding the gap junction protein Connexin-43^88^. *GJA1* was also found to be highly expressed in all NMM treated rDRG glial cultures (Fig. S4D).

To test if these pathways promote a pseudounipolar morphology (Fig. 5D), we first knocked down expression of *SEMA6A* in the rSGs using LV-shRNA transduction and puromycin selection. Selected glia were allowed to repopulate and then hiSNs were plated for terminal differentiation in coculture with shRNA-Sema6a expressing glia (shSema6a-rSGs) or shRNA-GFP expressing glia (shGFP-rSGs, control). Cells were cocultured for one week and then fixed and stained to visualize neuronal morphology (TUJ1, cytoskeletal marker, red) and glial expression of Sema6a (Sema6a, magenta; GFP, green) (Fig. 5E). We found that the iSNs differentiated in coculture with shSema6a-rSGs transition to pseudounipolar morphology at a reduced rate compared to controls (Fig. 5F, Table 6), with significantly fewer PU(T) cells (PUT *adj. residual*=-4.772, PU *adj. residual*=-6.861) compared to controls (PUT *adj. residual*=4.772, PU *adj. residual*=6.861), and significantly more multipolar cells (*adj. residual*=9.39) compared to controls (Multipolar *adj. residual*=-9.39. These results indicate that Sema6a-PlexinA4 signaling contributes to pseudounipolarization in this coculture model and may also decrease the incidence of multipolar morphology by inhibiting axonal outgrowth. However, the results also suggest that plexin signaling is not the sole mechanism underlying iSN maturation, since PU(T) cells make up only 10.28% of iSNs differentiated in shSema6a-rSG coculture.

**Table 6.** Raw frequency of morphologies observed in shRNA-Sema6a-SG coculture and shRNA-GFP-SG Coculture. X^2^ frequency statistic calculated for major morphology groups: Immature, Multipolar, PUT and PU. X^2^=94.62, df=3, p<0.001.

|  | <u>Morphology Frequencies</u> |  |  |  |  |  |  |  |  |
| --- | --- | --- | --- | --- | --- | --- | --- | --- | --- |
|  | Immature |  | Multipolar | Pseudounipolar Transition (PUT) |  |  | Pseudounipolar (PU) |  |  |
| Condition | Unipolar | Bipolar | Multipolar | PUT-Bulge | PUT-Bell | PUT-AB | PU-Short | PU-Long | Totals |
| shRNA-Sema6a | 2 | 9 | 242 | 15 | 7 | 2 | 4 | 1 | 282 |
| ShRNA-GFP | 20 | 8 | 181 | 38 | 29 | 14 | 53 | 16 | 359 |
| **Totals** | 39 |  | 423 | 105 |  |  | 74 |  | 641 |

To assess the role of satellite glial gap junctions in iSN maturation, we knocked down *GJA1* in the rSGs and plated iSNs for terminal differentiation in coculture with shRNA-GJA1 expressing glia (shGJA1-rSGs) or shRNA-GFP expressing glia (shGFP-rSGs, control) for one week (Fig 5D). Cocultures of hiSNs with shRNA-GJA1 expressing glia (shGJA1-rSGs) or shRNA-GFP expressing glia (shGFP-rSGs, control) were stained to visualize neuronal morphology (TUJ1, cytoskeletal marker, red) and glial expression of GJA1 (GJA1, magenta; GFP, green) (Fig 5G). This experiment was carried out in parallel with the semaphorin knockdown, and therefore we utilized a single group of sh-GFP RNA control samples. Strikingly, iSNs differentiated in coculture with shGJA1-rSGs also transition to pseudounipolar morphology at a reduced rate compared to controls (Fig. 5F, Table 7) with fewer PU(T) cells than expected (PUT *adj. residual*=-5.537, PU *adj. residual*=-7.014), and significantly more multipolar cells than expected (*adj. residual*=10.109). These results suggest that both gap junction and Sema6a/PlexinA mediated signaling contribute to pseudounipolarization.

**Table 7.** Raw frequency of morphologies observed in shGJA1-rSG coculture and shGFP-rSG Coculture. X^2^ frequency statistic calculated for major morphology groups: Immature, Multipolar, PUT and PU. X^2^=107.5, df=3, p<0.001.

|  | <u>Morphology Frequencies</u> |  |  |  |  |  |  |  |  |
| --- | --- | --- | --- | --- | --- | --- | --- | --- | --- |
|  | Immature |  | Multipolar | Pseudounipolar Transition (PUT) |  |  | Pseudounipolar (PU) |  |  |
| Condition | Unipolar | Bipolar | Multipolar | PUT-Bulge | PUT-Bell | PUT-AB | PU-Short | PU-Long | Totals |
| shRNA-GJA1 | 3 | 9 | 277 | 12 | 7 | 4 | 6 | 1 | 319 |
| ShRNA-GFP | 20 | 8 | 181 | 38 | 29 | 14 | 53 | 16 | 359 |
| Totals | 40 |  | 458 | 104 |  |  | 76 |  | 678 |

### Cocultured iSNs provide a more physiologically relevant model of neuropathy

iPSC derived neurons often exhibit embryonic phenotypes, which reduces the ability to model complex molecular mechanisms and disease processes ^9–15^. To determine whether cocultured iSNs provide a physiologically relevant model of peripheral neuropathy compared to iSNs differentiated alone, we examined the response of these cells to a common form of Chemotherapy Induced Peripheral Neuropathy (CIPN), paclitaxel induced peripheral neuropathy^145,146^. Patients with CIPN experience symptoms of allodynia, paresthesia, numbness, and pain in a stocking glove distribution, due to the degeneration of sensory nerve axon terminals^147^. We assessed the degeneration response of iSNs differentiated in coculture and alone when treated with low dose paclitaxel. At one week of terminal differentiation cultures were treated with paclitaxel chemotherapy or DMSO for 48 hours, then fixed and stained to visualize the neuronal cytoskeleton (TUJ1, Green) and gap junctions in both neurons and glia (GJA1, Magenta) (Fig. 6A,C). Neurodegeneration was estimated by quantifying TUJ1 signal.

**Figure 6.**
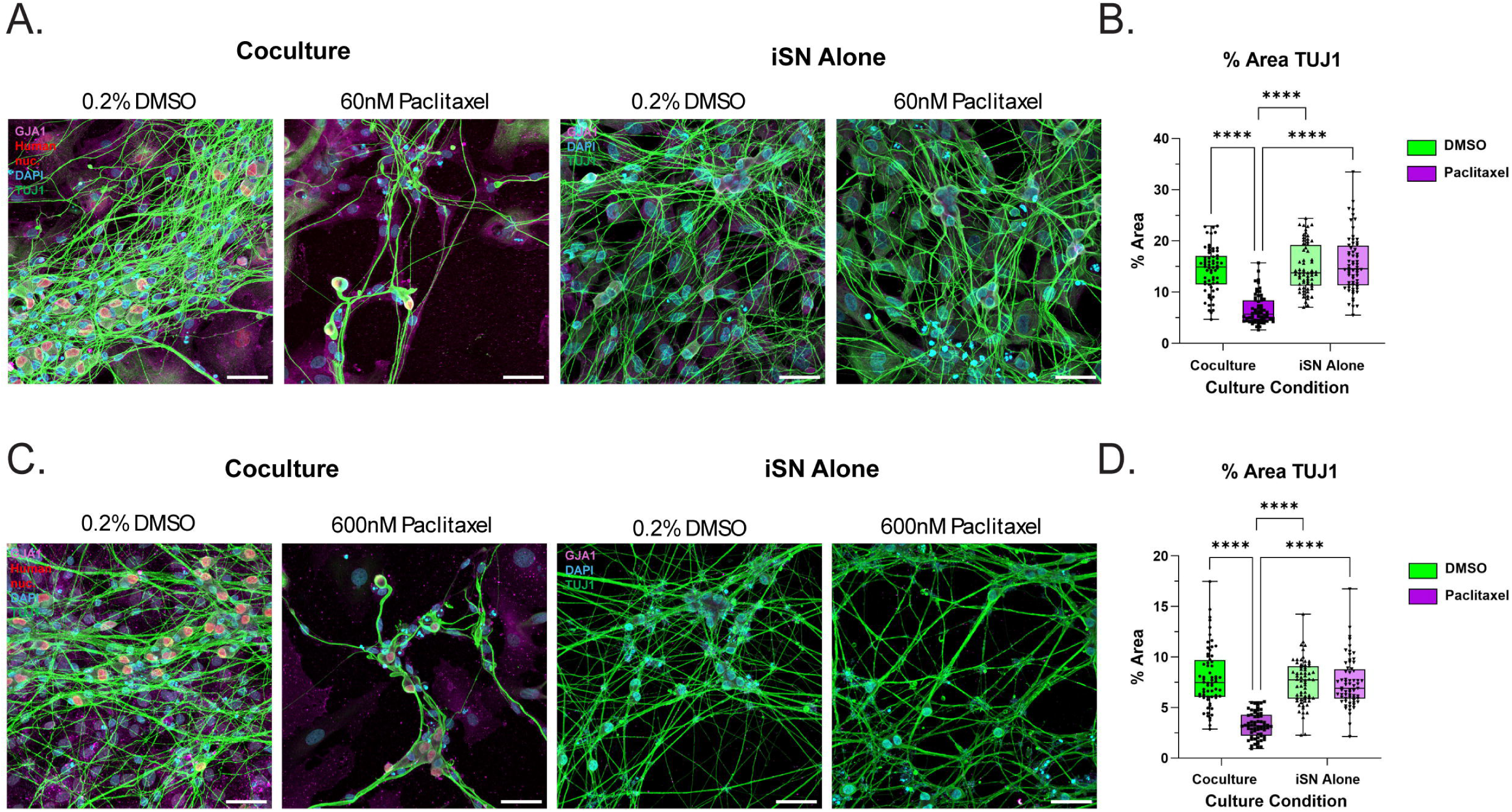
Cocultured iSNs are more susceptible to paclitaxel mediated neurodegeneration. A) Representative photomicrographs of Deng et al. 2023 generated iSNs in coculture or alone, treated with either 0.2% DMSO or 60nM paclitaxel. TUJ1= Green, GJA1=Magenta, Human Nuclei= Red, DAPI= Cyan. iSNs alone were not stained for human nuclei. Scale= 50um. B) Percent area occupied by TUJ1 particles in cocultured iSNs and iSNs differentiated alone + 0.2% DMSO or 60nM Paclitaxel. Two-Way ANOVA with Bonferroni post-hoc calculated in Graphpad Prism 10, p<0.05= significant, p<0.0001=****. n=64 fields from 4 biological replicates. C) Representative photomicrographs of Anatomic RealDRG iSNs in coculture or alone, treated with either 0.2% DMSO or 600nM paclitaxel. TUJ1= Green, GJA1=Magenta, Human Nuclei= Red, DAPI= Cyan. iSNs alone were not stained for human nuclei. Scale= 50um. D) Percent area occupied by TUJ1 particles in cocultured iSNs and iSNs differentiated alone + 0.2% DMSO or 600nM Paclitaxel. Two-Way ANOVA with Bonferroni post-hoc calculated in Graphpad Prism 10, p<0.05= significant, p<0.0001=****. n=64 fields from 4 biological replicates.

We first tested the impact of low-dose paclitaxel on the degeneration of iSNs differentiated with the Deng et al. 2023^15^ protocol, generated by NCATS. Analysis of TUJ1 expression showed that cocultured iSNs were significantly more susceptible to paclitaxel induced degeneration than iSNs differentiated alone (Bonferroni post-hoc: Coculture + Paclitaxel treatment vs. Coculture + DMSO: p <0.0001; vs. iSN Alone + Paclitaxel: <0.0001; vs. iSN Alone + DMSO: p<0.0001), which were not significantly impacted by treatment (Bonferroni post-hoc: iSN Alone + Paclitaxel vs. iSN Alone + DMSO p >0.9999) (Fig. 6B). We next tested the impact of the chemotherapy treatment on degeneration of RealDRG^TM^ iSNs from Anatomic, which require a larger dose of paclitaxel to elicit axonal degeneration^148^. Again, iSNs differentiated in coculture were more susceptible to paclitaxel induced degeneration compared to iSNs differentiated alone (Bonferroni post hoc: Coculture + Paclitaxel treatment vs. Coculture + DMSO: p <0.0001; vs. iSN Alone + Paclitaxel: <0.0001; vs. iSN Alone + DMSO: p<0.0001), which were not significantly impacted by treatment (iSN Alone + Paclitaxel vs. iSN Alone + DMSO p >0.9999) (Fig. 6D). Together these data indicate that iSNs cocultured with satellite glia provide a more physiologic model for toxin induced neuropathies.

## Discussion

The use of differentiated iPSCs to model human diseases has revolutionized our ability to understand pathology and develop efficacious treatments. While researchers have begun to utilize induced sensory neurons to understand peripheral neuropathies, current differentiation methods produce iSNs with an embryonic phenotype that do not fully recapitulate postnatal functions. In this study, we developed a protocol that enables maturation of iSNs in coculture with rodent satellite glia. We found that contact between iSNs and rSGs, mediated by semaphorin-plexin and glial gap junction signaling, promotes an adult morphology and physiology in the sensory neurons. Ultimately, this system also enables an enhanced model of neuropathy *in vitro*, by increasing iSN susceptibility to chemotherapy induced axonal degeneration at physiologic doses.

These studies establish a role for satellite glia in morphologic development of DRG neurons and demonstrate that we can leverage this relationship to reliably produce pseudounipolar iSNs. Previous studies have implicated DRG supporting cells in the morphologic maturation of sensory neurons, with mixed primary cultures of DRG neurons and glia facilitating the transition of sensory neurons to a pseudounipolar morphology^34–36,40^. This result has been recapitulated in induced neuro-mesodermal assembloids^149^, however iSNs cocultured with iPSC-derived^16^ or primary ^21^ Schwann cells alone do not exhibit advanced morphology. Based on these studies, we tested the possibility that satellite glia primarily contribute to sensory neuron maturation. *In vivo,* DRG sensory neurons increase physical contact with satellite glia during pseudounipolarization through the development of perikaryal outgrowths that increase contact surface area^40^ (Fig. S1B), In the current study, we show that physical contact between glial cells and iSNs is essential to the development of a pseudounipolar morphology, and RNA sequencing and immunohistochemistry indicate that specifically satellite glia are responsible for this effect.

Though glial contribution to sensory neuron maturation have been recognized previously, the molecular mechanisms enabling development of pseudounipolar morphology have been unclear. Since our results ruled out glial secreted factors as a direct mechanism of pseudounipolarization, we explored potential contact mediated mechanisms driven by our RNA sequencing data. In iSNs, we observed high expression levels of *Plexina4*, a receptor that interacts with glial semaphorins Sema3a and Sema6a^141–144,150^, which have both been implicated in sensory neuron axon development^141–144,151–153^. Since class 3 semaphorins are secreted, we assessed the role of Sema6a, a transmembrane semaphorin^154^, in the development of iSN axons. We found knockdown of glial *Sema6a* significantly reduced pseudounipolar formation in cocultured iSNs and instead resulted in a high proportion of multipolar cells. While there was a significant reduction in the proportion of pseudounipolar cells following *Sema6a* knockdown in the satellite glia, this effect was not complete. Factors contributing to observed low levels of pseudounipolarization may include incomplete knockdown of *Sema6a*, compensation via secreted Sema3a, and the possibility that semaphorin-plexin signaling is not the only mechanism underlying this process.

In the DRG, satellite glia communicate with adjacent satellite glia and sensory neurons through gap junctions^155,156^, contributing to the maintenance of neuronal homeostasis through direct exchange of diverse signaling molecules^157,158^. While little is known regarding gap junction signaling in sensory neuron axon development, communication through gap junctions correlates with neural crest cell differentiation into neuronal cells^159^. Knockdown of glial *GJA1,* like knockdown of Sema6a, significantly reduced the ability of glial cells to induce a transition of iSNs to a pseudounipolar morphology. As was seen following knockdown of Sema6a, knockdown of *GJA1* did not completely abolish pseudounipolarization. Again, this may be explained by incomplete knockdown efficiency, compensation by an alternate connexin, or another parallel signaling mechanism. Future studies need to be carried out to determine whether glial-glial or glial neuronal gap junction signaling contributes to development of pseudounipolar morphology.

RNA sequencing also revealed potential intracellular mediators of pseudounipolarization. Multiple molecules implicated in axonal guidance were significantly upregulated in cocultured iSNs, as illustrated in Table 2. These include *CDKL5,* a kinase linked to axonal polarization^98^ and *BCL11a,* a transcription factor that suppresses growth of supernumerary neurites and regulates microtubule protein expression^112^. We also observed significant downregulation of *RGMB* which promotes formation of multiple neurites^160^. The identification of cues and molecular pathways that stimulate pseudounipolar morphology may enable further refinements of iSN protocols.

Pseudounipolar morphology has multiple implications for sensory neuron physiology, as the axon stem may act as an axon initial segment that potentiates spontaneous activity^36^ or as a filter, gating nociceptive action potentials^161,162^. Importantly, we demonstrate that terminal differentiation in coculture yields more physiologically mature iSNs. Using MEA, we found that iSNs in coculture become electrically active earlier than iSNs cultured alone, and fire at a rate more consistent with rodent DRG neurons^54,55^. In addition to the development of an axon stem, the observed differential expression of several ion channels and receptors may contribute to mature firing rates. Upregulation of *SCN10a*, encoding Nav1.8, is of particular importance, as this sodium channel maintains repetitive action potential firing in DRG nociceptors^163^ and is integral to pain signaling^72,73,164^. Increased expression of muscarinic receptors *CHRM2* and *CHRM3* may also contribute to spontaneous activity, as acetylcholine has been implicated in the development of bursting and spiking behavior in sympathetic neurons cocultured with satellite glia^165^. Lastly, increased neuronal excitability may reflect a reduction in expression of several potassium channels, namely *KNCT2*^166^ and *KCNS1*^167^, which have previously been linked to hyperexcitability and neuropathic pain.

In this study our goal was to generate iSNs with a more mature adult-like phenotype that would allow us to better model peripheral neuropathies, and we tested the efficacy of modeling toxic neuropathy using chemotherapy treated cocultured iSNs. We found that cocultured iSNs were significantly more susceptible to paclitaxel chemotherapy mediated axonal degeneration compared to iSNs alone. Hyperexcitability of sensory neurons has been shown to contribute to axonal degeneration across multiple neuropathies^6^, including chemotherapy induced peripheral neuropathy (CIPN)^145,146^, and so the increased baseline excitability of cocultured iSNs may enhance susceptibility to degeneration. Satellite glia may also directly contribute to neuronal degeneration, as studies in rodent models of CIPN indicate that glial release of inflammatory cytokines contribute to degeneration^168–170^. Overall, our results provide new avenues for studying the potential involvement of satellite glial-sensory neuron interactions in the pathogenesis of neuropathies, as this protocol can easily be used to model diabetic peripheral neuropathy, inherited neuropathies, or axonal injury as well.

In summary, our work identifies molecular direct contact cues whereby satellite glia support sensory neuron development and establishes an accessible protocol for the reliable differentiation of mature iSNs which can be used to study physiological and pathological changes in sensory neurons and develop novel treatments for peripheral neuropathies.

*Limitations of this Study:*

This study is limited by the use of rodent glial cells and human derived sensory neurons. Currently, there are no publicly available protocols to differentiate human satellite glia, however several protocols have been developed to induce Schwann Cells^38,171^ which share a glial progenitor cell lineage. Adjustments of these protocols such that satellite glia differentiation can be optimized will allow for an advanced iSN differentiation protocol with exclusive use of human cells, and aid in identification of molecular mechanisms underlying normal human sensory neuron development and neuropathy.

## Methods

### Animal Care

Sprague-Dawley rats were purchased from Charles River Laboratory. Animals were cared for in accordance with NIH standards and guidelines. Experimentation was approved by the Dana Farber Cancer Institute IACUC.

### iPSC Culture

Induced pluripotent stem cells (iPSCs) from LiPSC-GR1.1 (RRID:CVCL_RL65, Male, Hispanic, normal, <1D, umbilical cord blood, Lonza Inc. via NIH RMP, MCB p28) and NCRM5 cell lines (RRID: CVCL_1E75, Male, normal, <1D, umbilical cord blood, Lonza Inc. via NIH RMP, MCB p28) were cultured in feeder-free conditions in Essential-8 Medium (Thermo Fisher; A1517001) using cell culture plates or flasks coated with vitronectin (Thermo Fisher; A14700). iPSCs were passaged with EDTA (Thermo Fisher; 15575020) every 3 days at a 1:6 ratio. Cells were treated with the CEPT cocktail for 24 hours after passaging (Chroman 1, MedChem; HY-15392; Emricasan, SelleckChem S7775, Polyamine supplement (1000X), Sigma-Aldrich P8483, Trans-ISRIB, R&D Systems 5284)^172^. Basic characterization including, authentication using G-banded karyotyping, and Mycoplasma detection using Mycoalert kit (Lonza; LT07-318) were carried out at p28. Pluripotency was assessed at p30 using the TaqMan hPSC Scorecard assay (Thermo Fisher; A15876) and analysis software, and significance was determined using the *Student’s t-test*. iPSCs (p30) were differentiated into neural crest precursors (NCPs) and immature sensory neurons (SNs) using our published protocol^15^.

### rDRG-iSN Coculture

#### Rodent DRG glial cell culture

Sprauge-Dawley day E14.5 rat DRGs were dissected. DRGs were trypsinized (1mg trypsin /mL HBSS) for 50 minutes at 37°C. 40k cells/well were plated on uncoated Lab-Tek™ II Chambered Coverglass (Thermofisher cat#: 155409) with filtered Glia Base Media (DMEM/F12, 10% FBS, 1% Pen. Strep., 0.08% glucose). After a 30-minute incubation at 37°C 7.5% CO_2._, chambers were vigorously shaken, and media was fully changed to remove cells in suspension. Remaining glial cells were cultured in Glia Base Media at 37°C 7.5% CO_2_ for one week with media changes 3x/ week.

#### Terminal differentiation and iSN-rDRG glia co-culture

Frozen Neural Precursor Cells (NPCs) were differentiated by NIH-NCATS Stem Cell Laboratory using the protocol outlined in Deng et al., 2023^15^. Lab-Tek™ II Chambered Coverglass (Thermofisher cat#: 155409) were coated with 0.005% poly-ornithine solution (Sigma-Alrich cat#: P4957) in dH2O 1 hr at 37°C, washed, and then coated with 5 ug/mL mouse laminin (Thermofisher cat#: 23017015) in dH2O 1 hr at 37°C. NPCs were thawed in NCATS Maturation Media (DMEM/F12+Glutamax, 2% B27,1% N2, 25ng rh-BDNF/b-NGF/GDNF/NT3) + CEPT rock inhibitor cocktail (50nM Chroman-1 (Medchem Express Cat#: HY-15392), 5uM Emricasan (Selleck cat#: S7775), 0.001% Polyamine supplement (Sigma-Alrich cat#: P8483), 0.7uM Trans-ISRIB (Tocris cat#: 5284)). 15k cells/well were plated in NCATS Maturation Media + CEPT on coated cultureware for terminal differentiation alone (iSN-Only), or in coculture with rDRG glial cultures treated with NCATS Maturation Media + CEPT. 24 hrs after plating, media was changed to NCATS Maturation Media + 1uM PD0332991 (Tocris cat#: 4786). Cells were cultured for one week at 37°C 5% CO_2._, with ½ media changes 3x/week.

For each experiment throughout this study, we indicate the number of biological replicates used, with each biological sample representing an experiment done with one set of iSNs that were plated and tested individually. The experiment was subsequently replicated on additional sets of cells plated and tested on a different day (the biological replicates). n=4 biological replicates for experiments performed in Figure 1C, n= 2 NCRM5 and n=2 GR1.1.

### Quantification of iSN morphology frequency

#### Immunofluorescence

Cells were fixed for downstream Immunofluorescence (IF) on Day 14. First, cells were fixed with 2% paraformaldehyde (PFA) (EMS cat#: 15710; 1:1 4% PFA to spent media) 10 minutes at room temperature (RT), and then in 4% PFA 20 minutes at RT. Cells were washed 3x in PBS and stored at 4°C. Fixed cultures were permeabilized with 0.1% Triton-x in PBS 10 minutes and blocked in blocking buffer (5% Normal Goat Serum (Cell Signaling Tech. cat#: 5425S) in 1x PBS) for 1 hr at RT. Cultures were then incubated overnight at 4°C in primary antibodies (Table 11) diluted in blocking buffer. The following day, cultures were washed and incubated in Alexa-Fluor secondary antibodies (Table 12) and DAPI diluted 1:1000 in blocking buffer for 1 hr at RT. Co-Cultures were stained with mouse anti-human nuclei-CY3 conjugate (Sigma cat#: MAB1281C) for 2 hrs at RT to distinguish human neuronal nuclei from rodent. Cells cultured in Lab-Tek™ II Chambered Coverglass were stored at 4°C in PBS prior to imaging, and cells cultured on coverslips were mounted on slides with fluoromount (Thermofisher cat#: OB10001).

**Table 11.**
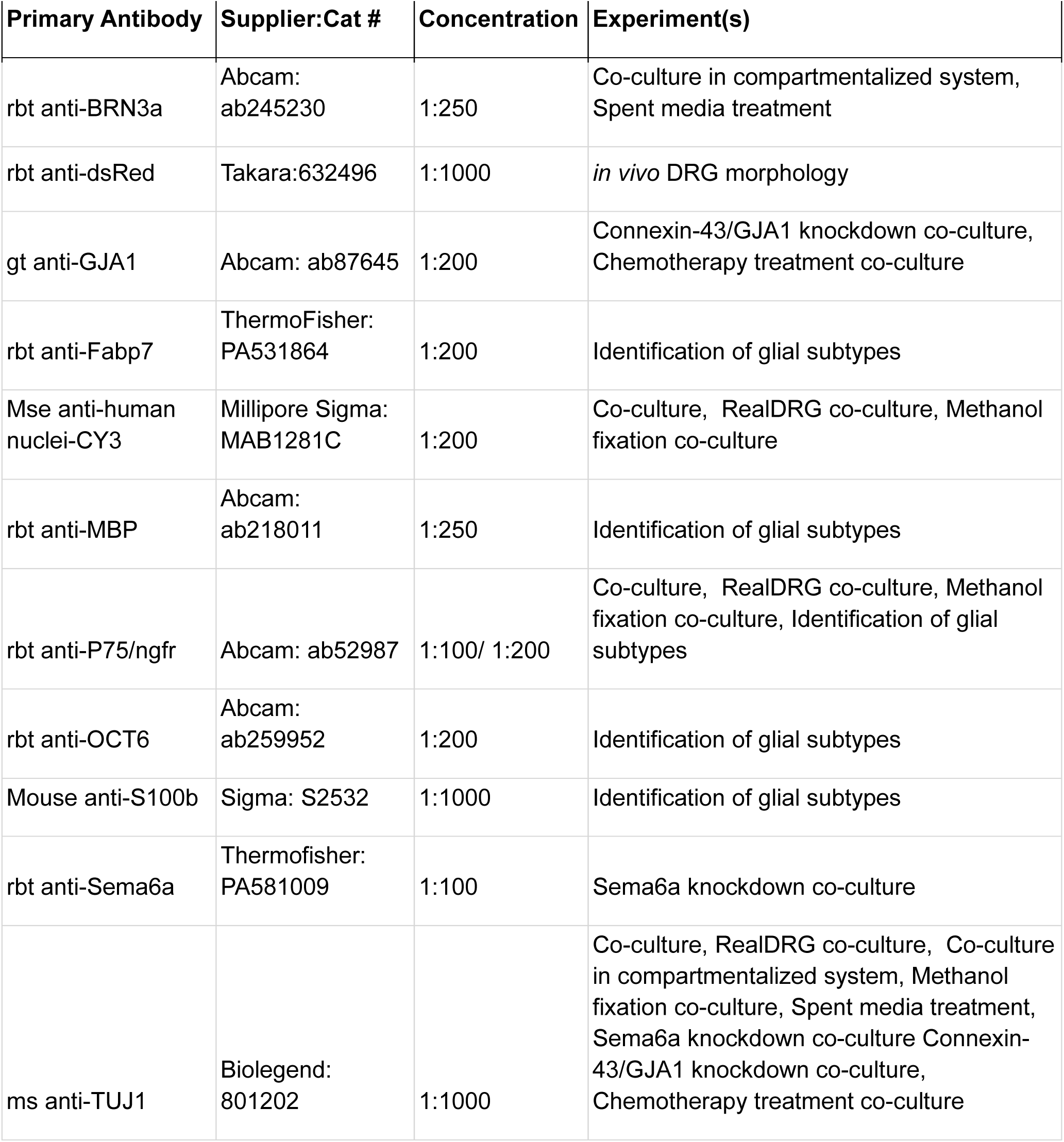
Primary antibodies.

**Table 12.**
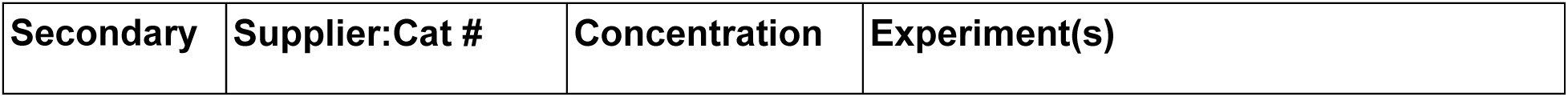
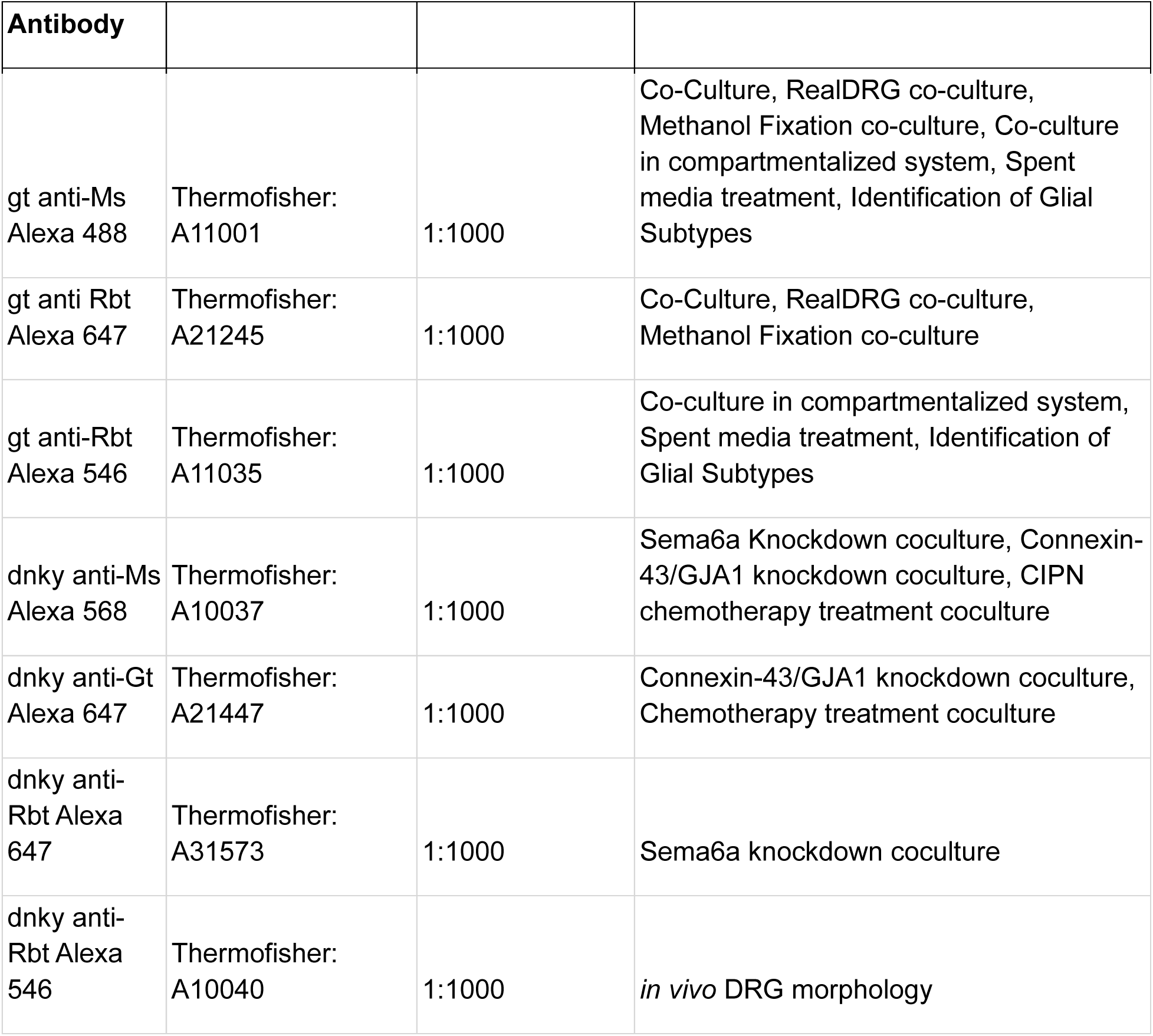
Secondary antibodies.

| Secondary | Supplier:Cat # | Concentration | Experiment(s) |
| --- | --- | --- | --- |

| Antibody |  |  |  |
| --- | --- | --- | --- |
| gt anti-Ms<br>Alexa 488 | Thermofisher:<br>A11001 | 1:1000 | Co-Culture, RealDRG co-culture,<br>Methanol Fixation co-culture, Co-culture<br>in compartmentalized system, Spent<br>media treatment, Identification of Glial<br>Subtypes |
| gt anti Rbt<br>Alexa 647 | Thermofisher:<br>A21245 | 1:1000 | Co-Culture, RealDRG co-culture,<br>Methanol Fixation co-culture |
| gt anti-Rbt<br>Alexa 546 | Thermofisher:<br>A11035 | 1:1000 | Co-culture in compartmentalized system,<br>Spent media treatment, Identification of<br>Glial Subtypes |
| dnky anti-Ms<br>Alexa 568 | Thermofisher:<br>A10037 | 1:1000 | Sema6a Knockdown coculture, Connexin-43/GJA1 knockdown coculture, CIPN<br>chemotherapy treatment coculture |
| dnky anti-Gt<br>Alexa 647 | Thermofisher:<br>A21447 | 1:1000 | Connexin-43/GJA1 knockdown coculture,<br>Chemotherapy treatment coculture |
| dnky anti-Rbt Alexa<br>647 | Thermofisher:<br>A31573 | 1:1000 | Sema6a knockdown coculture |
| dnky anti-Rbt Alexa<br>546 | Thermofisher:<br>A10040 | 1:1000 | *in vivo* DRG morphology |

#### Imaging

All photomicrographs of cells cultured in Lab-Tek™ II Chambered Coverglass were collected on the Zeiss-980 airy-scan/confocal microscope. Images of all conditions were obtained at 20x with 1.3x zoom and bidirectional scanning in SRY-4 Airy-Scan. Multiplex imaging was utilized with 405, 488, 561, and 639 lasers at 800V gain, 1% laser power, and 0 offset. All images were obtained in a z-stack with 0.35um step size. Systematic random sampling was utilized to image n=50-100 cells/biological replicate across 8-wells. Areas with fused cell bodies and axonal fasciculations were omitted from sampling. All images were 3D airy-scan post processed in Zen Blue software.

#### Analysis

All human cells in each photomicrograph were categorized into the following morphologic groups : unipolar (one axonal projection off of the cell body, that does not bifurcate), bipolar (two axonal projections off of opposite cell body poles, ∼180° angle), multipolar (more than two axonal projections), Pseudounipolar Transition (PUT)-Bulge (two axonal projections off of the bottom of the cell body, axons form >90° angle), PUT-Bell (two axonal projections off of the bottom of the cell body, axons form a 90° angle or less), PU-Associated Branch (PU-AB,T-shaped pseudounipolar axon with a branch <10uM long), PU-Short (T-shaped pseudounipolar axon, with an axon stem <60um long), PU-Long (T-shaped pseudounipolar axon, with an axon stem >60um long). Human cells were distinguished from rodent cells based on expression of human nuclei-Cy3 marker. Images were blinded and intra-rater or inter-rater reliability was calculated to be >90%. Morphology frequency was compared across morphologic groups: Immature (unipolar and bipolar), Multipolar, Pseudounipolar Transition (PUT-Bulge, PUT-Bell, PUT-AB), and Pseudounipolar (PU-Short and PU-Long) using the Chi-square test in Graphpad Prism 10. Sub-morphologies were pooled into major morphologic categories to reach chi-square test validity requirements. Adjusted residuals with Bonferroni correction were calculated in excel using standard formulas.

### Multi-Electrode Array Recording

#### Cell Culture

rDRG glia were dissected and plated in 48-well, 16 electrode Cytoview-MEA plates (Axion) at 20k cells/well in Glia Media. After a 30-minute incubation at 37°C 7.5% CO_2,_ plates were vigorously shaken, and media was fully changed to remove cells in suspension. Remaining glial cells were cultured in Glia Media at 37°C 7.5% CO_2_ for one week with media changes 3x/ week. On the 7^th^ day, iNCPs were thawed and plated in NCATS Maturation Media + CEPT at 150k-200k cells/ well, in coculture or alone on coated wells (100ug/mL PDL 1 hr 37°C, 10ug/mL mouse laminin 1 hr 37°C). A fraction of the wells remained glial cells alone as a control, treated with NCATS Maturation Media +CEPT. 24 hrs after plating, media was changed to NCATS Maturation Media + 1uM PD0332991. Cells were cultured for 5-8 weeks at 37°C 5% CO_2._, with ½ media changes 3x/week. n=4 biological replicates; replicates 1-3 = iPSC Gr1.1, replicate 4 = iPSC NCRM5.

#### MEA Recording

MEA recordings were performed on the Maestro Pro recording system from Axion at 37°C 5% CO_2_, 1x/week for 5 minutes, after at least 5 minutes of acclimation. AxIS navigator software was used to record standard Spontaneous Neural and Viability properties. Burst Detection was enabled with the following specifications: Single Electrode Minimum # Spikes=5, Network Minimum #Spikes=50, Maximum Interspike Interval= 100s, Mean Firing Rate detection window= 100s. Recordings were carried out weekly, and data was analyzed when an average of 7+ active electrodes was reached for both cocultured iSNs and iSNs alone. Electrodes were considered active with a minimum of 5 spikes/minute.

#### Analysis

Data was analyzed at the time point when an average of 7+ electrodes were active in both conditions. Wells with 2 or less active electrodes were excluded from analysis. Glial cells were not active, and therefore were excluded from all analyses aside from # of active electrodes. Data did not follow a normal gaussian distribution, and therefore averages were compared using the Mann-Whitney U test in Graphpad Prism 10. n=# wells, pooled from 4 biological replicates.

### RNA sequencing of iSN and rDRG Glia cocultures

#### Cell Culture and RNA Collection

rDRG glia were dissected and plated in uncoated glass bottom 35mm Fluorodishes (WPI cat #: FD35-100) at 250k cells/dish in Glia Media. After 30 minutes at 37°C 7.5% CO_2,_ dishes were vigorously shaken, and media was fully changed to remove cells in suspension. Remaining glial cells were cultured in Glia Media at 37°C 7.5% CO_2_ for one week with media changes 3x/ week. On Day 7, cells were either treated with NCATS Maturation Media + 1uM PD0332991, maintained in Glia Media, or co-cultured with iSNs as described above. iSNs were plated at 90k/dish in co-culture or alone, as described above. Cultures were maintained at 37°C 5% CO_2._, with ½ media changes 3x/week for an additional week. On Day 14, RNA was collected from Glia Media Treated rDRG Glia, NCATS Maturation Media Treated rDRG Glia, iSN-rDRG Glia Co-Culture, and iSN-Only mass cultures using the RNeasy Plus Mini Kit (Qiagen cat#: 74134). n=3-4 biological replicates; Coculture: n= 2 iPSC NCRM5, n=1 GR1.1. iSN only: n=4 iPSC NCRM5. Glial cells derived from n=4 rats.

#### RNA sequencing

mRNA bulk sequencing was carried out at the Dana Farber Cancer Institute Molecular Biology Core Facility. The genomics core team performed RNA QC, rRNA depletion, library preparation (Roche Kapa mRNA Kit), and sequencing on the Illumina NovaSeq 6000. Reads were aligned to both the rat (*Rattus norvegicus*) and human genome using STAR aligner to separate rDRG glia and iSN reads in co-culture. GEO Accession #: GSE272868.

#### Analysis

Normalized counts (with raw counts <3 set to NA) were analyzed for differential gene expression using the DESeq2 R package version 1.42.0^173^ with an adjusted p-value cutoff of 0.05. Gene Ontology Enrichment Analysis was performed using the clusterProfiler R package version 4.10.0^174^ with an adjusted p-value cutoff of 0.05.

### iSN-rSG co-culture in transwell compartmentalized system

#### Cell Culture

rDRG glia were dissected as described above and plated at 40k cells/well on uncoated Millicell® Standing Cell Culture Inserts (MilliporeSigma cat#: PIHP01250). rDRG glia were prepared as described above. On the 7^th^ day, iNPCs were plated at 15k cells/ well on sterile 12mm #1.5 coverslips, coated as described above, in the same plate for terminal differentiation either in compartmentalized co-culture or alone. Cells were allowed to attach to coverglass for 30 min at 37°C 5% CO_2_, after which Glia Media was removed from rDRG glial inserts and inserts were transferred to wells with NPCs dedicated for terminal differentiation in compartmentalized co-culture. 24 hrs after iNPC plating, media was changed to NCATS Maturation Media + 1uM PD0332991. Cells were cultured for one week at 37°C 5% CO_2._, with ½ media changes 3x/week. On the 14^th^ day, rDRG glial culture inserts were moved to new wells and glia attached to insert were visualized with CellTracker Red CMTPX (Thermofisher cat#: C34552). Cells were incubated with 12.5uM dye in DMEM/F12 for 30 min at 37deg 5% CO2, after which media was changed to DMEM/F12 and live cells were imaged at 10x TRITC on the Nikon-Ti E microscope. iSNs adherent to coverglass were fixed for downstream Immunofluorescence (IF) as described above. n=3 biological replicates, iPSC Gr1.1.

#### Imaging and Analysis

Photomicrographs of cells cultured on coverslips were collected on the Nikon Ni-E confocal microscope at 60x with oil immersion. Images were obtained with 405 and 488 lasers, at 30V gain 5% power and 561 30V gain 7% power. All images were obtained in a z-stack with 0.35um step size. Systematic random sampling was utilized to image n=50-100 cells/biological replicate across 3 coverslips. Areas with fused cell bodies and axonal fasciculations were omitted from sampling. Inter-rater reliability was calculated >90%. Morphology frequencies and statistics were carried out as described above.

### iSN cultures treated with co-culture spent media

Spent media was collected from iSN-rDRG Glia Co-Cultures over the course of one week and stored at 4°C. On Day 21, iNPCs were plated alone on chambered coverglass in NCATS Maturation Media + CEPT as described above. After 24 hrs, media was either changed to ½ fresh NCATS Maturation Media + 1uM PD0332991 + ½ spent media or maintained in NCATS maturation Media + 1uM PD0332991 as described above. Cells were cultured for one week with 3x full media changes maintaining specified ratios. On Day 28, cultures were fixed for IF and images were acquired and analyzed as described. n=3 biological replicates; n= 2 iPSC NCRM5 n= 1 iPSC Gr1.1.

### iSN-rSG co-culture with glial methanol fixation

rDRG glia were dissected, plated, prepared, and cultured in chambered coverglass for one week as described above. On Day 7, glia were treated with NCATS maturation media + CEPT, and 24 hrs later media was changed to NCATS Maturation Media + 1uM PD0332991. Glia were maintained in NCATS Maturation Media + 1uM PD0332991 an additional week with ½ media changes 3x/week. On day 14, cells were fixed in 95% methanol 7 min at 4°C, washed, and treated with NCATS maturation media + CEPT for downstream iSN co-culture. iNPCs were thawed and plated at 15k cells/well for terminal differentiation in iSN-rDRG Glial Co-Culture with methanol fixation, iSN-rDRG Glial Co-Culture without methanol fixation (prepared as described above), or iSN-only culture (prepared as described above). 24 hrs after iNPC plating, media was changed to NCATS Maturation Media + 1uM PD0332991. Cells were cultured for one week at 37°C 5% CO_2._, with ½ media changes 3x/week. After one week of coculture, cells were fixed with PFA for downstream IF and images and analyses were carried out as described above. n=4 biological replicates, n=2 NCRM5, n=2 GR1.1.

### iSN-rSG co-culture with glial *Sema6a* or *GJA1* sh-RNA knockdown

rDRG glia were dissected, plated, prepared, and cultured for five days as described above. On Day 5, Glia were transduced with ShRNA-Sema6a-puro-GFPtag lentivirus (Millipore Sigma Cat#: SHCLNV), ShRNA-GJA1-puro-GFPtag lentivirus (Millipore Sigma Cat#: SHCLNV), or ShRNA-GFP-puro control lentivirus (Millipore Sigma cat#: SHC005V) in Glia Media for 24 hrs, after which media was changed. Cells were maintained in Glia Media 72 hrs post-transduction and evaluated for sh-RNA expression using live-imaging in brightfield and FITC on the Nikon-Ti E microscope. Glia were then treated on Day 8 with 1ug/mL puromycin in Glia Media for 48hrs. Media was changed, and selected cells were allowed to proliferate for four days. On Day 14, iNPCs were plated at 15k cells/well in NCATS Maturation Media + CEPT for terminal differentiation in iSN-rDRG Glia Knockdown Co-Culture, iSN rDRG Glia Knockdown Control Co-culturel and iSN-rDRG Glia Co-Culture Control (prepared as described above). 24 hrs after iNPC plating, media was changed to NCATS Maturation Media + 1uM PD0332991, and cells were co-cultured for one week at 37°C 5% CO_2._, with ½ media changes 3x/week. On Day 21, co-cultures were fixed for IF as described. n=4 biological replicates, 2/ iPSC line.

### Modeling chemotherapy induced axonal degeneration

#### Chemotherapy treatment of iSN co-culture with rDRG glia

rDRG glia were prepared as described above. iSNs generated with the Deng et al. 2023^15^ protocol, were plated at 15k cells/well in rDRG glial co-culture or alone and maintained in culture for 7 days as described above. On day 14, cells were treated with 60nM Paclitaxel (Sigma Aldrich cat #: T7402) or 0.2% DMSO (Sigma Aldrich cat#: D2650) in NCATS Maturation Media + 1uM PD0332991 for 48 hrs. n=4, iPSC NCRM5. Anatomic RealDRG iSNs were plated at 5k cells/well in rDRG glial co-culture or alone and maintained in culture for 7 days as described above. On day 14, cells were treated with 600nM Paclitaxel (Sigma Aldrich cat #: T7402) or 0.2% DMSO (Sigma Aldrich cat#: D2650) in NCATS Maturation Media + 1uM PD0332991 for 48 hrs. n=4, Anatomic RealDRGs are derived from one genetic iPSC line. Cells were then fixed as described above for IF.

#### Immunofluorescence (IF)

Fixed cultures were permeabilized with 0.1% Triton-x in PBS 10 minutes, and blocked in blocking buffer (5% Normal Donkey Serum (Sigma Aldrich cat#: D9663) in PBS) for 1 hr at RT. Next, they were incubated overnight at 4°C in primary antibodies (Table 11) diluted in blocking buffer. The following day, cultures were washed and incubated in Alexa-Fluor secondary antibodies (Table 12) and DAPI diluted 1:1000 in blocking buffer for 1 hr at RT. Cells were washed and stored in PBS at 4°C prior to imaging.

#### Imaging

All photomicrographs of cells cultured in Lab-Tek™ II Chambered Coverglass were collected on the Zeiss-980 airy-scan/confocal microscope. Images of all conditions were obtained at 20x with 1.3x zoom and bidirectional scanning in SRY-4 Airy-Scan configuration. Multiplex imaging was utilized with 405, 488, 561, lasers at 800V gain, 1% laser power, and 0 offset. All images were obtained in a z-stack with 0.35um step size. Systematic random sampling was utilized to image n=16 fields/biological replicate across 6-wells. All images were 3D airy-scan post processed in Zen Blue software.

#### Analysis

Particle analysis was carried out on gray-scale TUJ1 channel image stacks. Threshold was manually set at the brightest z-stack slice for each image, and percent area occupied by TUJ1 pixels was quantified for each image stack. Average values were calculated for each field. Field averages were compared between treatment groups and culture conditions using two-way ANOVA analysis with the Bonferroni post-hoc test in Graphpad Prism. n=64 fields from 4 biological replicates.

## Acknowledgements

NCI R01 CA205255. Schematics created with Biorender.

## Author Contributions

Conceptualization: CJL, OYT, RAS. Generation of NCPs: KG, VJ, DC, PO, CT, IS. Performance of experiments: CJL, MFPM, ES, SD, VP, SZ, OYT. Analysis: CJL, MFPM, MT, TM. Funding Acquisition: RAS. Project Administration: CJL, RAS. Supervision: RAS, CW. Writing: CJL, RAS, with input from all authors.

## Supplemental Methods

### Animal Care

Sprague-Dawley rats were purchased from Charles River Laboratory. Ntrk2-CreER/C57BL/6 mice and lox-tdtomato/C57BL/6 mice were shared by Dr. David Ginty, Harvard Medical School. PLP1-GFP/C57BL/6 mice were shared by Dr. Lisa Goodrich, Harvard Medical School. Animals were cared for in accordance with NIH standards and guidelines. Experimentation was approved by the Dana Farber Cancer Institute IACUC.

### Tamoxifen induced sparse labeling of Ntrk2+ sensory neurons with tdtomato *in vivo*

Ntrk-CrER mice were crossed with lox-tdtomato mice and subsequently crossed with PLP1-GFP/C57BL/6 mice. Timed pregnant Ntrk-CreER/lox-tdtomato/PLP-1GFP mice were weighed and tamoxifen (TMXF) dose was calculated at 40ug/g body weight. 2mg/mL Tamoxifen (TMXF) was stored in methanol at -80C and the final TMXF dose was diluted in 100uL corn oil. Diluted TMXF was vortexed 45 minutes and dried in a Speedvac to remove methanol from the mixture, leaving TMXF in corn-oil alone. Injections were carried out intraperitoneally at E12.5-E14.5 to sustain expression and sacrifice at E14.5, E16.5, E18.5, P5. At time of sacrifice, embryonic lumbar DRGs were dissected, and expression of tdtomato and PLP1-GFP was evaluated via epifluorescence. DRGs were then drop fixed in 4% paraformaldehyde at 4°C overnight. The next day, DRGs were washed 3x with 1XPBS and stored at 4°C in 1X PBS until staining. n=3+ embyros total per age group.

### Quantification of DRG sensory neuron morphology frequency *in vivo*

#### Immunofluorescence

Fixed DRGs were permeabilized 2 hrs in 0.5% tritonX in PBS, and then blocked in 5% Normal Donkey Serum (NDS)/0.5% tritonX in PBS 4-5hrs at RT. They were then incubated in rbt anti-dsRed (Takara Cat#: 632496) primary antibody diluted 1:1000 in blocking buffer at 4°C overnight, and washed the next day with 1X PBS. Next, DRGs were incubated with dnky anti-rbt 546 (Thermofisher Cat#: A10040) secondary antibody diluted 1:500 in blocking buffer for 3 hrs at RT. After staining, DRGs were stored at 4°C in 1XPBS, or gradually transferred to methanol for tissue clearing.

#### DRG Tissue Clearing

Stained DRGs stored in PBS were gradually transferred to methanol with 10 minute incubations in the following concentrations: 25%, 50%, 75%, and 100%. After methanol dehydration, each DRG was transferred to a slide and placed within a slide spacer (VWR Cat#102096-614). BABB solution (1 Benzyl Alcohol:2 Benzyl Benzoate) was added to the DRG and incubated 2-3min, after which it was replaced with fresh BABB, and steps were repeated until the tissue was cleared. Then, a circular coverslip was mounted on top with tissue embedded in BABB solution.

#### Imaging

Confocal images were acquired on the Nikon Ni-E confocal microscope, first at 20X to visualize the entire DRG, and then at 40x to visualize individually labeled neurons. Images were acquired at x resolution, 2.4 speed, 0.2-0.3um step, with 488 and 561 laser at 1.04 laser power, 36 gain, 0 offset. All cells labeled with tdtomato in the DRG were imaged from the top to bottom of neuronal cell bodies.

#### Analysis

Images were blinded after acquisition, and NIS elements software was used to visualize all cell morphology in 2D and/or 3D. All labeled cells were categorized into the following morphologic groups: bipolar (elongated cell body, with an axon emanating from the two opposite poles of the cell body with ∼180° angle between), intermediate (round cell body with two axons forming <180° angle between), and pseudounipolar (one axon stem emanating from cell body, forming a T-Shape axon). Cells were considered apoptotic with condensed cell bodies small, and no axons attached.

### Anatomic RealDRG iSN co-culture with rDRG glia

Frozen RealDRG iSNs were purchased from Anatomic (cat#: 1020). rDRG glia were dissected, plated, prepared, and cultured for one week as described above. On Day 7, RealDRG iSNs were thawed in Chrono-Senso-MM (Anatomic) and plated at 15k, 9k, or 5k cells/well either in rDRG glia co-culture in NCATS Maturation Media. Lab-Tek™ II Chambered Coverglass were coated with 0.01% Poly-ornithine solution for 1 hr at 37°C, washed with dPBS, and coated with iMatrix-511 (Reprocell cat#:NP892-021) 1:50 in dPBS 3 hrs at 37°C for iSN-only culture control. After 24 hrs, media was either changed to NCATS maturation Media + 1uM PD0332991. Cells were cultured for one week with 3x ½ media changes. On Day 14, cells were fixed for downstream IF and analysis as described in text. n=3 biological replicates at densities indicated.

### Treatment of rDRG glia with neuronal maturation media

rDRG glia were dissected, plated, prepared, and cultured for one week as described above. On Day 7, glia were either treated with NCATS Maturation Media + CEPT and cultured at 37°C 5% CO_2_ or maintained in Glia Media at 37°C 7.5% CO_2_. After 24 hrs, NCATS Maturation Media + CEPT was changed to NCATS Maturation Media + 1uM PD0332991. Glia were matured for an additional week with ½ media changes 3x/week and fixed for downstream Immunofluorescence (IF) on Day 14. First, cells were fixed with 2% paraformaldehyde (PFA) (EMS cat#: 15710; 1:1 4% PFA to spent media) 10 minutes at room temperature (RT), and then in 4% PFA 20 minutes at RT. Cells were washed 3x in PBS and stored at 4°C for downstream experimentation. n=4 biological replicates.

### Identification of Glial Subtypes in rDRG Glial cultures with IF

#### Immunofluorescence

Fixed rDRG glial cultures treated with and without NCATS maturation medium were stained using the protocol described in text.

#### Imaging

All photomicrographs of cells cultured in Lab-Tek™ II Chambered Coverglass were collected on the Zeiss-980 airy-scan/confocal microscope. Images of all conditions were obtained at 20x with 1.3x zoom and bidirectional scanning in SRY-4 Airy-Scan configuration. Multiplex imaging was utilized with 405, 488, 561, lasers. All images were obtained in a z-stack with 0.35um step size. Systematic random sampling was utilized to image n=12-15 fields/biological replicate across 8-wells. All images were 2D airy-scan post processed in Zen Blue software. n=4 biological replicates.

#### Analysis

Particle analysis was carried out on glial marker channels. Threshold was manually set at brightest z-stack slice for each image, and percent area occupied by particles, average particle size, and average mean gray value were quantified for each image stack. Average values were calculated for each field, and averages were compared between treatment groups using the t-test function in excel where n=# of fields.

### Quantification of rSG-iSN contacts in coculture with methanol fixation

#### Immunofluorescence and Imaging

IF and imaging were performed as described in text. n=4 biological replicates

#### Analysis

Number of rSGs contacting individual iSNs was quantified using the CellCounter plugin in FIJI. Contact was defined as apposition between P75+, human Nuc. Cy3-processes with TUJ1+, Human Nuc. Cy3+ cell body and neurites. Number of rSGs were marked by individual DAPI+, Human Nuc. Cy3-nuclei and rSGs contacting iSNs without a nucleus present in the field were not counted. Contacts were counted per individual iSN, and each iSN was categorized into defined morphologic categories. Average number of total glial contacts/cell was compared using the t-test function and average number of total glial contacts/cell for each morphologic category was compared using Two-way ANOVA with Bonferroni post-hoc test in GraphPad Prism 10.

### Supplemental Figure Legends

**Figure S1.**
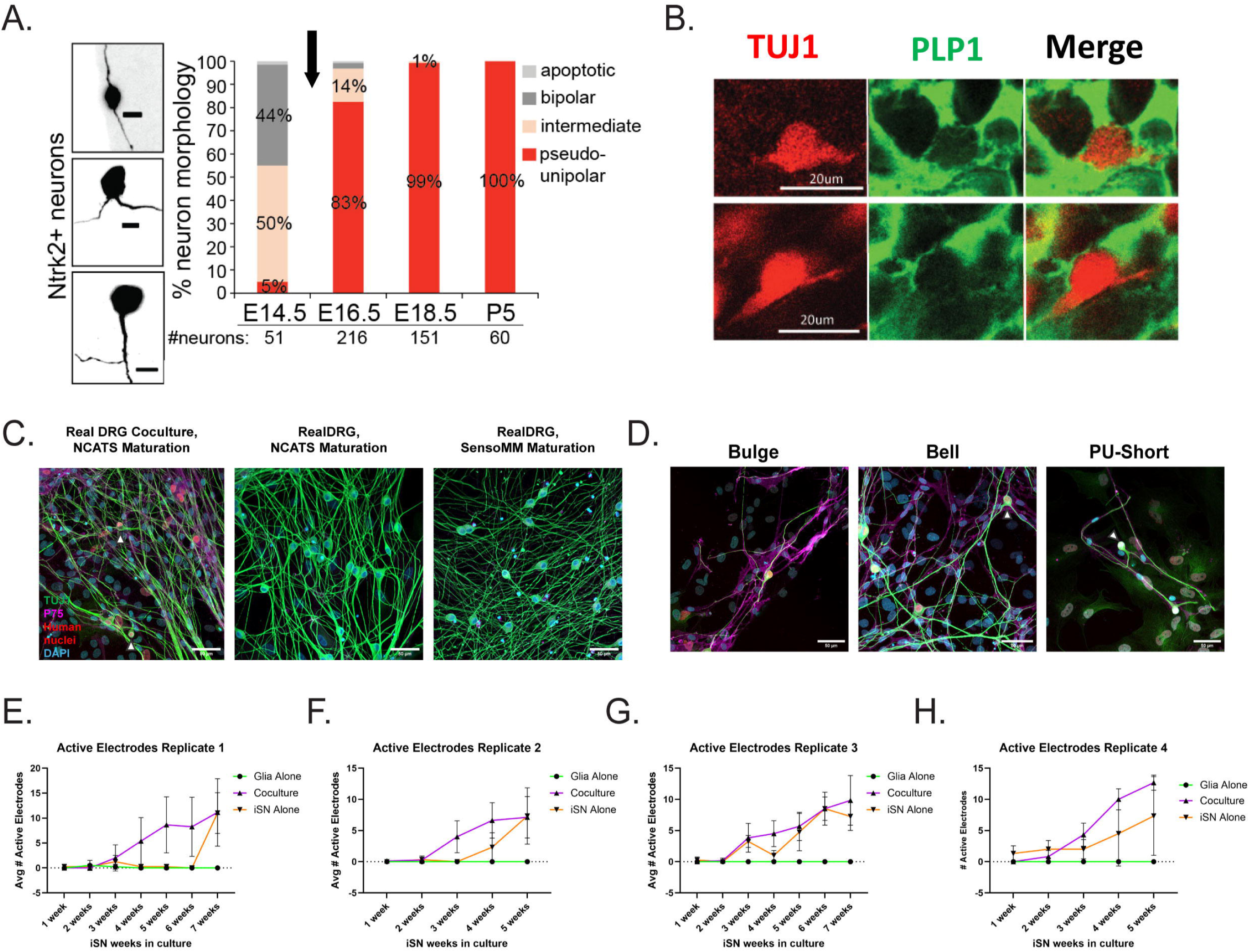
Human iSNs differentiated in coculture with embryonic rodent DRG glia exhibit advanced morphology and physiology. A) Timeline for pseudounipolar transition was established *in vivo* in embryonic mice expressing tamoxifen induced NTRK2-tdtomato. Morphology frequencies from whole DRGs were quantified and categorized as apoptotic (light gray), bipolar (dark gray), intermediate (pink), or pseudounipolar (red). DRG NTRK2+ neurons became pseudounipolar between days E14.5 and E16.5, indicated by black arrow. E14.5 3-5 DRGs were quantified. B) Photomicrographs illustrating physical contact between satellite glia and DRG neurons during pseudounipolarization. Green= PLP1, Red= Ntrk-tdtomato. Scale= 20um. C-D) C. Representative photomicrographs of Anatomic RealDRGs terminally differentiated in coculture or alone with NCATS maturation media and Anatomic’s proprietary media Senso-MM. D. Photomicrographs depicting transitional and pseudounipolar morphologies observed in cocultured Anatomic RealDRGs. TUJ1=Green, p75=Magenta, human nuclei = Red, DAPI=Cyan. RealDRGs alone were not stained for human nuclei. Scale=50um.E-H) Average number of active electrodes during weekly MEA recordings for fourbiological replicates. Cocultured iSNs=purple, iSNs alone= orange, Glia alone= green. Average >7 active electrodes were observed in both cocultured iSNs and iSNs alone at 7 weeks in replicate 1, 5 weeks in replicate 2, 6 weeks in replicate 3, and 5 weeks in replicate 4.

**Figure S2.**
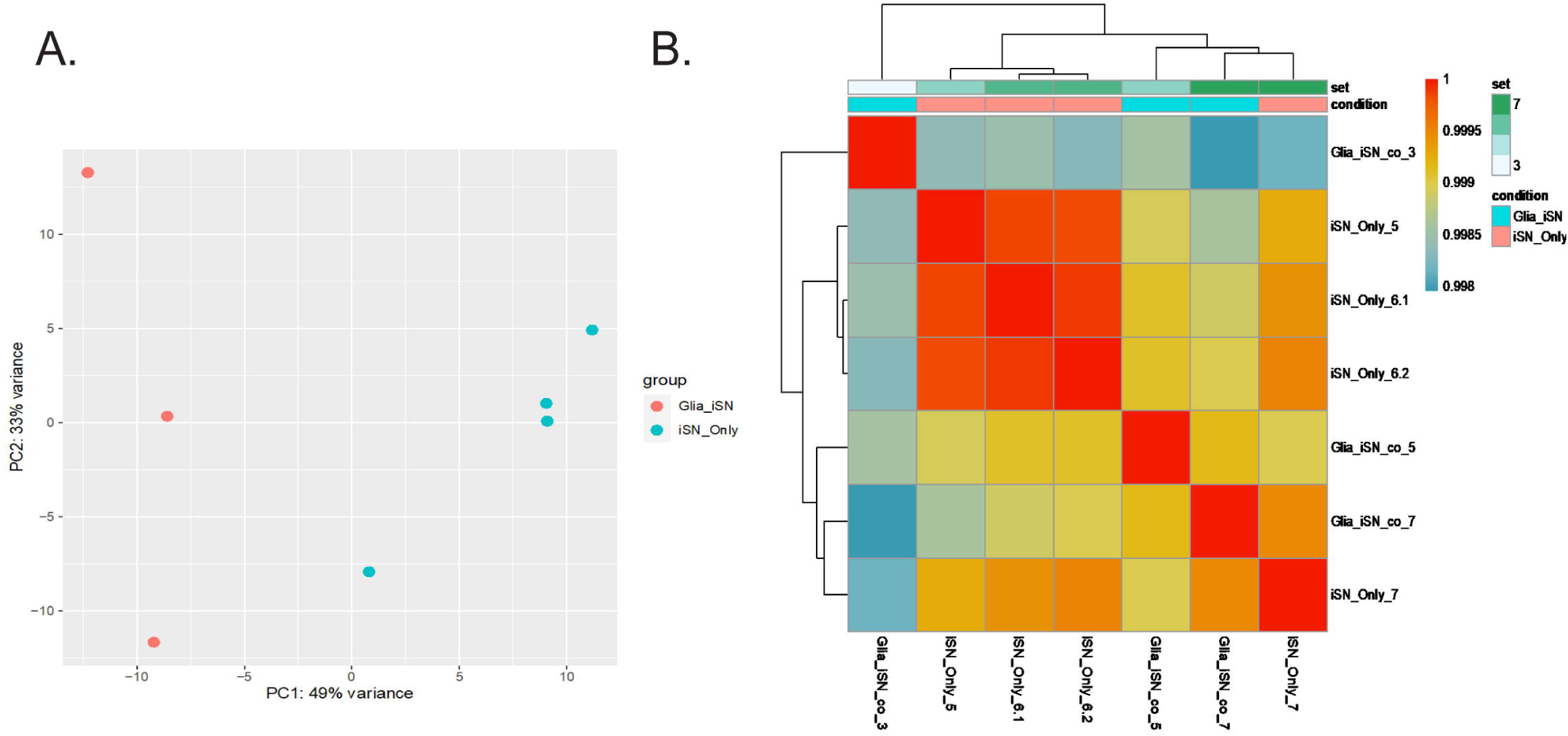
iNCPs terminally differentiated in co-culture more advanced in neuronal differentiation compared to iNCPs alone. A) Principal component analysis and B) Hierarchal Cluster analysis of RNA sequencing reads from cocultured iSN and iSNs alone.

**Figure S3.**
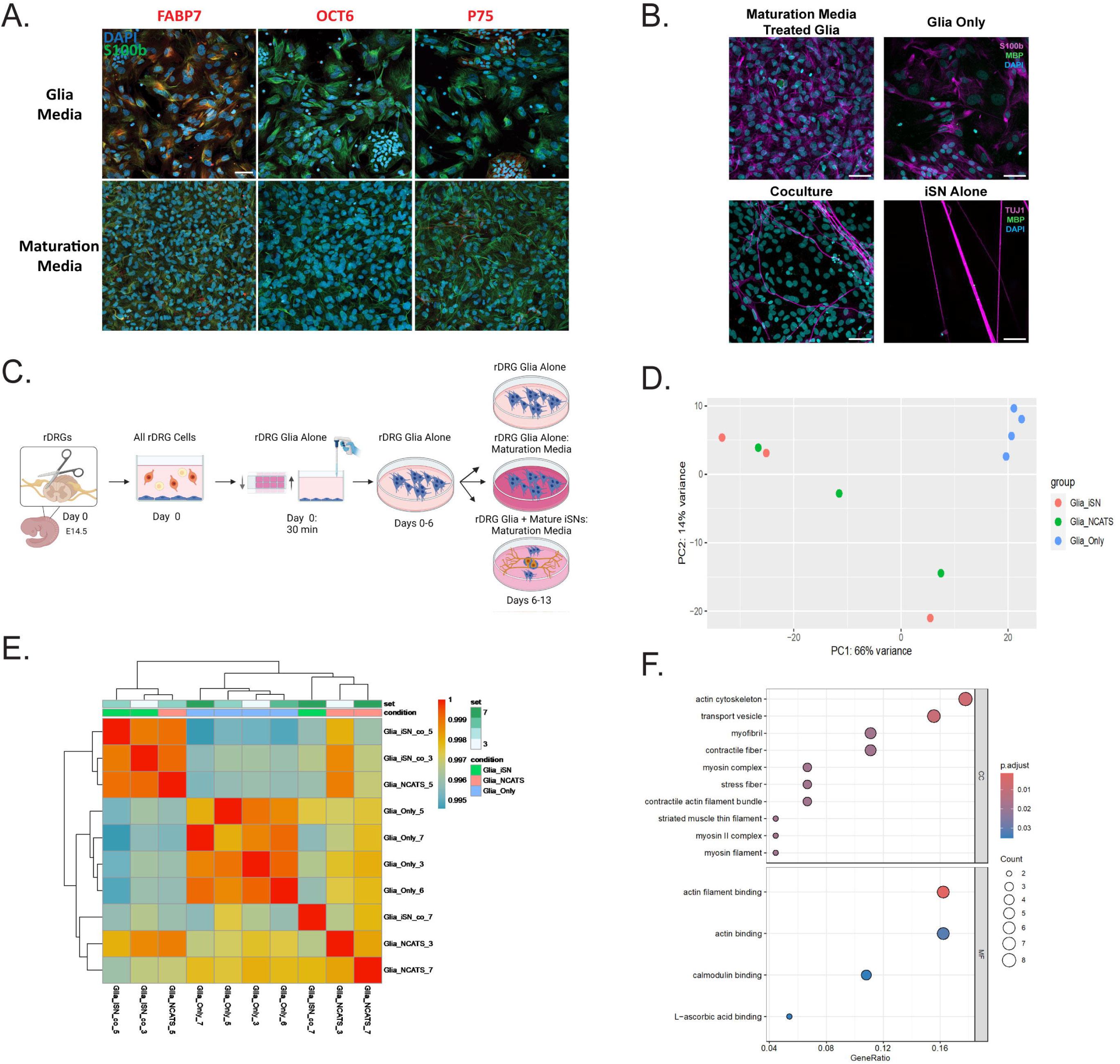
Figure 3. Rodent DRG satellite glia (rSGs) are responsible for advanced maturation of iSNs. A) Representative photomicrographs of glial cultures maintained in glial base media for 1 week and treated with NCATS Maturation Media for 1 week (Maturation Media) or maintained in glial base media for 2 weeks (Glia Media). S100b= Green, FABP7/OCT6/P75= Red, DAPI= Cyan. Scale = 50 um. B) Representative photomicrographs of glial cultures maintained in glial base media for 1 week and treated with NCATS Maturation Media for 1 week (Maturation Media) or maintained in glial base media for 2 weeks (Glia Only), iSNs in coculture and iSNs alone. Glial Cultures: S100b=Magenta, MBP=Green, DAPI=Cyan, iSN cultures: TUJ1=Magenta, MBP=Green, DAPI= Cyan. No MBP signal was detected in any culture condition. Scale=50um. C) Schematic of glial culture protocol for RNA sequencing. D-E) Principal component analysis (D) and Hierarchal Cluster analysis (E) of RNA sequencing reads from rDRG glia in coculture, treated with maturation media, and maintained in base glia media. F) GO terms significantly downregulated in cocultured glia compared to glia treated with maturation media. P adj.<0.05.

**Figure S4.**
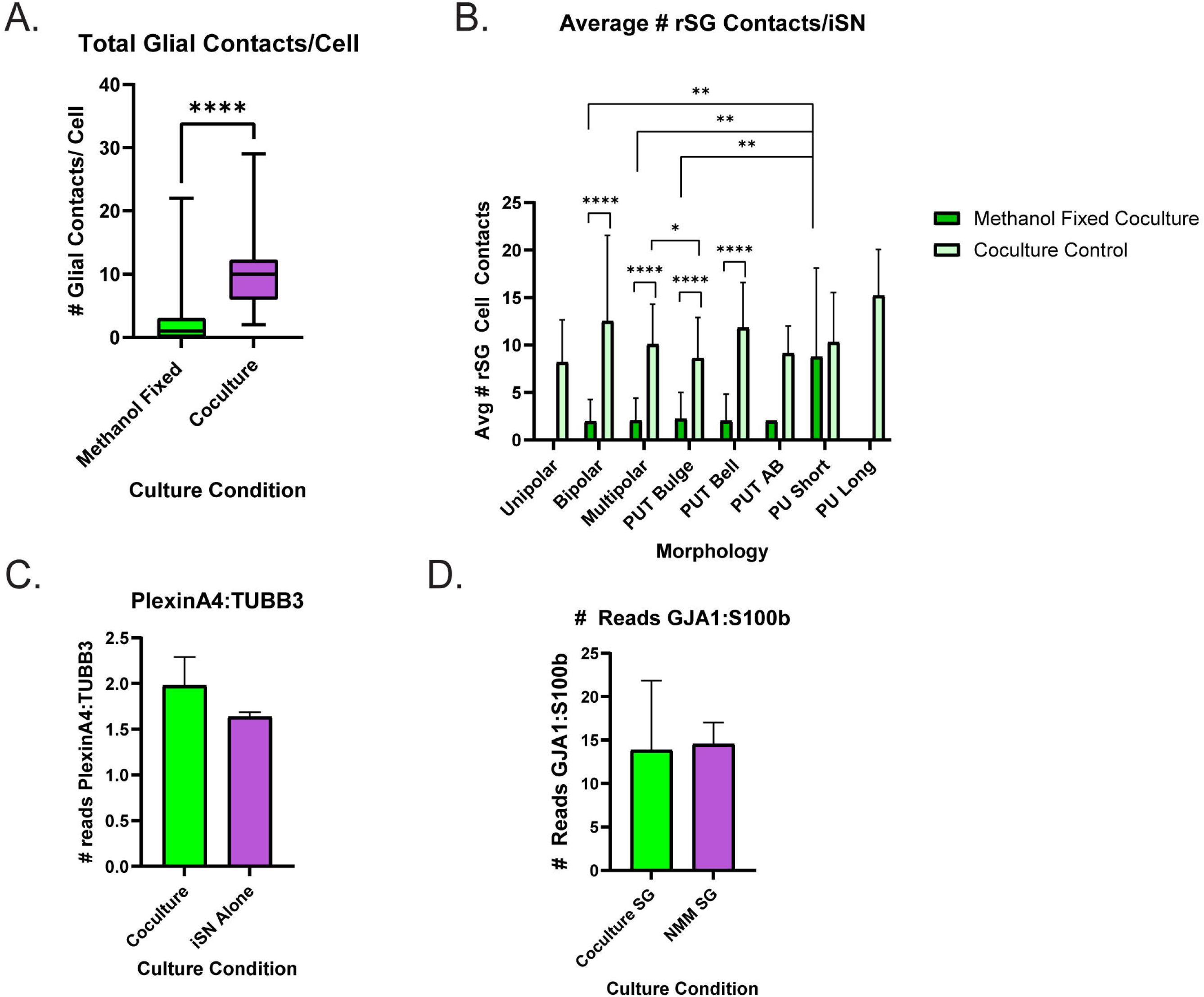
Physical contact between rSGs and hiSNs is essential to pseudounipolarization. A) Total rSG-iSN contacts per individual iSN in coculture with methanol fixed glia or in normal coculture. T-test performed in GraphPad Prism 10, ****=p<0.0001. B) Average number of rSG-iSN contacts per individual iSN in coculture with methanol fixed glia or in normal coculture, partitioned by morphology of iSN. Two-way ANOVA with Bonferroni post-hoc test calculated in GraphPad Prism 10. P<0.05=*, P<0.01=**, P< 0.0001=****. C) PlexinA4 mRNA reads normalized to TUBB3 (TUJ1) mRNA reads in cocultured iSNs and iSNs differentiated alone. D) GJA1 mRNA reads normalized to S100b mRNA reads in cocultured and maturation media treated glia.

### Supplemental Index

1. Document S1: Supplemental Methods and Figure Legends
2. Document S2: Supplemental Figures 1-4
3. Supplemental Tables Containing Large Datasets:

a. S1: Cocultured iSN vs iSN Alone DGE
b. S2: Cocultured iSN vs iSN Alone GO
c. S3: NMM treated rDRG Glia vs rDRG Glia alone in base DGE
d. S4: NMM treated rDRG Glia vs rDRG Glia alone in base GO
e. S5: Cocultured rDRG Glia vs rDRG Glia Alone in base DGE
f. S6: Cocultured rDRG Glia vs rDRG Glia alone in base GO
g. S7: Cocultured rDRG Glia vs NMM treated rDRG Glia DGE
h. S8: Cocultured rDRG Glia vs NMM Treated rDRG Glia GO

